# Boltz2-Notebook: An Interactive Google Colab Platform for Diffusion-Based Biomolecular Structure and Binding Affinity Prediction using the Boltz2 model

**DOI:** 10.64898/2026.09.18.752645

**Authors:** Atharva Tilewale, Dhaval Patel

**Affiliations:** Department of Industrial Biotechnology, Gujarat Biotechnology University, Near Gujarat International Finance Tec (GIFT)-City, Gandhinagar, 382355, Gujarat, India

**Keywords:** protein-ligand structure prediction, binding affinity prediction, Boltz-2, Google Colab, accessible bioinformatics software, biomolecular foundation models, BindingDB benchmark, reproducible computational workflows

## Abstract

Recent advances in deep learning-based structure prediction, including AlphaFold3 and the open-source Boltz model family, have extended biomolecular modeling to joint prediction of protein-ligand, protein-nucleic acid, and multi-chain complexes with binding-affinity estimation. Boltz-2 is among the most feature-complete of these open models, but its practical use requires a local CUDA-capable GPU, command-line execution, and manually authored YAML configuration files, limiting accessibility for researchers without dedicated computational infrastructure. We developed Boltz2-Notebook, a Colab-native interface comprising four integrated stages - automated environment setup, interactive parameter-to-YAML generation, execution management, and automated confidence and affinity visualization - together with a manifest-driven batch mode for multi-target screening. All modelling capabilities are inherited unmodified from Boltz-2; Boltz2-Notebook’s contributions are limited to accessibility, input construction, and workflow automation. Independent of the software, we curated a benchmark of 317 protein-ligand pairs (122 proteins, 277 ligands) from BindingDB and predicted binding affinity in triplicate using the Boltz-2 command-line engine on high-performance computing infrastructure. Predicted and experimental pIC50 values showed moderate correlation (Pearson r = 0.609 [95% CI 0.540-0.675]; Spearman ρ = 0.625; R^2^ = 0.371; MAE = 0.968 pIC50 units), with high triplicate reproducibility (pairwise r = 0.97) but a systematic compression of the predicted affinity range and no measurable relationship between Boltz-2’s self-reported confidence metrics and prediction accuracy. Boltz2-Notebook is freely available as open-source software and provides external, reproducible evidence - including a specific confidence-calibration limitation - relevant to interpreting Boltz-2 affinity predictions responsibly.

## 1. Introduction

Predicting the three-dimensional structure of proteins and their complexes from sequence has been one of the central open problems in computational biology for decades. The introduction of AlphaFold2 (Jumper et al., 2021) represented a step change in this field, achieving near-experimental accuracy for single-chain protein structure prediction and rapidly displacing physics-based and homology-based methods as the default approach for structural inference. Its success catalyzed a wave of follow-on work extending deep learning-based structure prediction to protein complexes, protein-nucleic acid assemblies, and, most recently, protein-small-molecule interactions. AlphaFold3 (Abramson et al., 2024) extended this paradigm to a unified diffusion-based architecture capable of jointly modeling proteins, DNA, RNA, ligands, ions, and covalent modifications within a single complex, establishing the template for what are now commonly termed biomolecular "foundation models" - general-purpose structure predictors not restricted to a single molecular class. This shift has been significant for drug discovery and structural biology alike, because protein-ligand interactions, not protein structure in isolation, are what ultimately determine most therapeutic and functional outcomes.

A parallel, open-source line of development has pursued the same goal outside of proprietary or access-restricted release models. Boltz-1 (Wohlwend et al., 2024) reimplemented and extended the AlphaFold3-style co-folding architecture as a fully open, freely redistributable model, explicitly framed around democratizing access to biomolecular structure prediction. Boltz-2 (Passaro et al., 2025) built on this foundation to address a limitation that neither AlphaFold3 nor Boltz-1 had resolved: while both models substantially improved complex structure prediction, neither reliably predicted binding affinity, a property that is often of greater practical interest than pose accuracy alone for hit identification and lead optimization. Boltz-2 reports strong correlation with experimental affinity across multiple benchmark contexts, including performance approaching that of free-energy perturbation methods on the FEP+ benchmark (Wang et al., 2015) at markedly lower computational cost, and outperforming submitted methods on the CASP16 affinity assessment track. Together with its structure-conditioning features - including template integration, user-defined distance constraints, and support for covalent, cyclic, and modified-residue chemistry - Boltz-2 offers one of the most feature-complete open biomolecular modeling toolkits currently available.

But there are real-world constraints on who can take advantage of this capability. To run Boltz-2 as released requires a local NVIDIA GPU with a working CUDA installation, a python environment with a specific and version sensitive set of dependencies, and command line execution of the prediction pipeline. Inputs are to be specified as hand-authored YAML files, according to a schema encoding entity types (protein, DNA, RNA, ligand), optional per-residue modifications, and constraint blocks (covalent bonds, binding pockets, contact restraints, templates) - a specification that is expressive, but unforgiving of syntactic or semantic errors, and that assumes familiarity with both its internals and standard bioinformatics file formats. In addition to installation and input preparation, a complete analysis pipeline also requires downstream scripting for extracting and interpreting confidence outputs (pLDDT, predicted aligned error, affinity scores), creating structural visualisations, and managing results across multiple jobs, none of which are included with the base package. These requirements are a real, well-documented barrier to entry for researchers who lack access to local high-performance computing, do not have a CUDA-capable workstation or substantial command-line and scripting experience, and disproportionately affect smaller academic groups, teaching contexts, and researchers whose primary expertise is experimental rather than computational.

This barrier is not new to the field, and prior tools have addressed analogous versions of it for earlier-generation models. ColabFold (Mirdita et al., 2022) removed the local-installation requirement from AlphaFold2 and RoseTTAFold by combining an accelerated MMseqs2-based homology search (Steinegger & Söding, 2017) with a Google Colab-hosted execution environment, enabling structure prediction from a browser with no local GPU or dependency management, and has since become one of the most widely used structural bioinformatics tools as a result. More recently, DeepMind released the AlphaFold Server, a free, browser-accessible interface to AlphaFold3 for non-commercial use. However, the AlphaFold Server restricts inputs to a curated, non-custom ligand library, does not support binding-affinity prediction, imposes daily job-count limits for academic users, and explicitly prohibits automated or high-throughput screening-style use in its terms of service (Google, 2024) - constraints that make it unsuitable for exploratory small-molecule work or batch analysis. Critically, neither ColabFold nor the AlphaFold Server addresses Boltz-2 specifically: no equivalent zero-install, browser-based interface currently exists that exposes Boltz-2’s full input schema - custom small-molecule ligands, multi-entity DNA/RNA complexes, user-defined constraints, and affinity prediction - without requiring local installation or manual YAML authoring.

We developed Boltz2-Notebook to close this specific, narrow gap: an accessible, cloud-native, end-to-end interface to Boltz-2 built entirely on the free tier of Google Colab. The platform is intended neither to modify nor to extend Boltz-2’s underlying modeling capability, but to make that capability usable without local GPU infrastructure, CUDA configuration, or hand-written YAML, by providing guided input construction, automated execution, and integrated result visualization and analysis within a single notebook-based workflow. In this sense, Boltz2-Notebook occupies the same relationship to Boltz-2 that ColabFold occupies to AlphaFold2: an accessibility layer, not a modeling contribution.

Separately from the accessibility question, the practical reliability of Boltz-2’s affinity predictions outside of the benchmarks reported in its original description remains comparatively unexamined. Independent, external evaluation on data assembled after a model’s release is an important complement to developer-reported benchmarks, both because it tests generalization to differently curated data and because it allows explicit assessment of properties, such as the reliability of self-reported confidence metrics, that are not always characterized in the original publication. BindingDB (Liu et al., 2007) provides a large, continually updated, literature-curated repository of experimentally measured protein-ligand binding affinities well suited to this purpose. We therefore used a newly curated, quality-filtered subset of BindingDB to independently evaluate Boltz-2’s affinity prediction performance, alongside the development of Boltz2-Notebook itself.

The objective of this work is twofold and the two components are presented as distinct contributions. First, we describe the design and implementation of Boltz2-Notebook, an interactive, Colab-native platform that provides guided parameter construction, automated execution, and integrated confidence and affinity analysis for Boltz-2, without requiring local GPU hardware, CUDA installation, or manual YAML configuration. Second, we report an independent evaluation of Boltz-2’s protein-ligand binding affinity prediction performance on a curated, single-chain BindingDB benchmark, including an assessment of the relationship between the model’s self-reported confidence metrics and its actual predictive accuracy. Together, these contributions are intended to lower the practical barrier to using Boltz-2 and to provide external evidence relevant to its use in affinity-driven research contexts.

## 2. Implementation and Software Architecture

### 2.1 Overall Software Architecture

Boltz2-Notebook is organized as a sequential, four-stage pipeline implemented across a small set of purpose-specific Python modules (setup.py, param_gen.py, Boltz_Run.py, analysis.py), each corresponding to one cell of a Google Colab notebook (Figure 1). This modular separation mirrors the natural stages of a structure-prediction workflow - environment preparation, input specification, model execution, and results interpretation - and allows each stage to be executed, inspected, or re-run independently within the same notebook session. Three notebook variants share this architecture: a stable single-job notebook (V1) supporting protein-ligand and multi-chain prediction with affinity analysis; an advanced notebook (V2) that extends the same pipeline with the additional entity types and constraint mechanisms described in §2.3; and a batch notebook (§2.6) that replaces the single-job execution and analysis stages with a manifest-driven engine for processing many targets in one run. Table 1 summarizes the feature set available in each of the three notebook variants.

**Figure 1.**
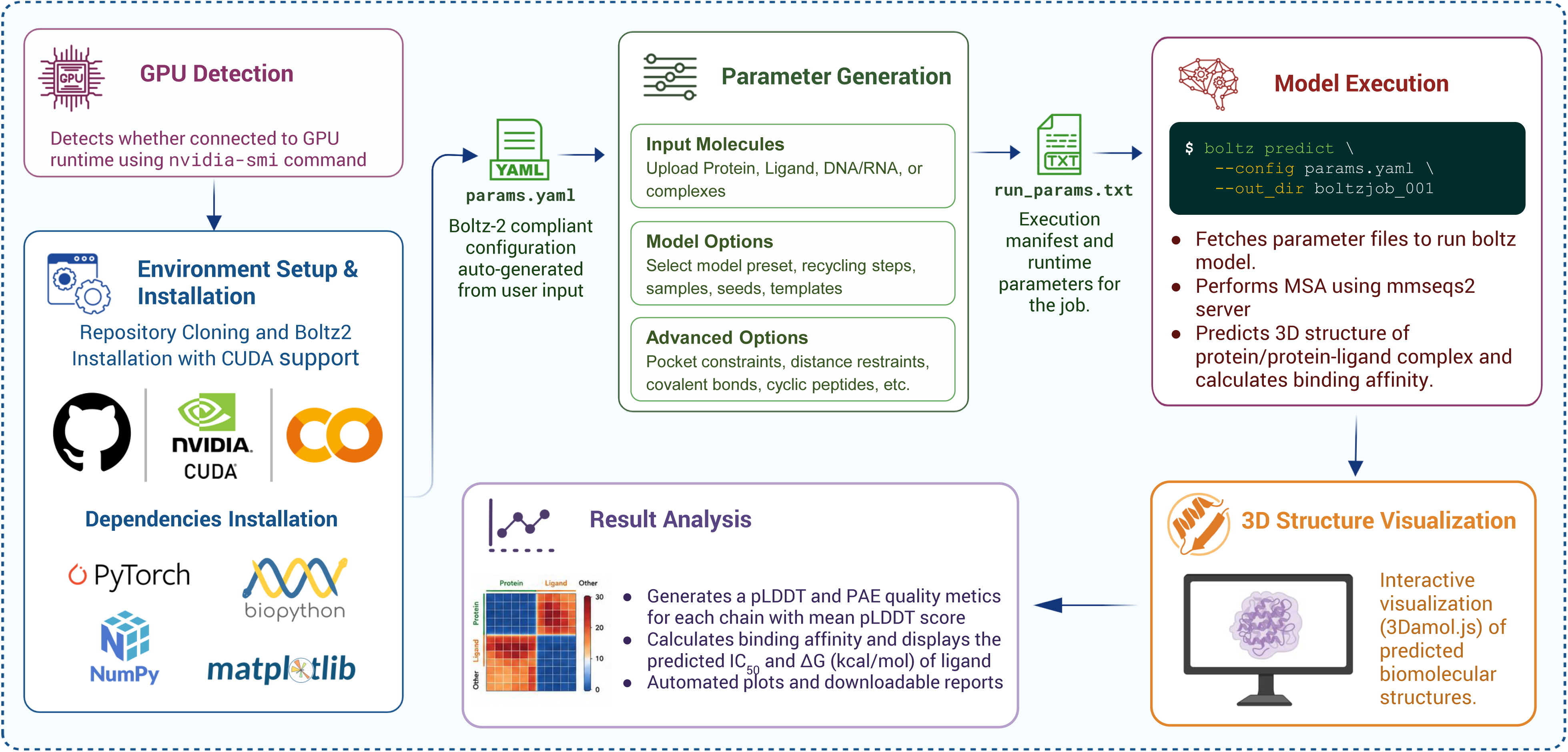
Overall architecture and data flow of Boltz2-Notebook. Schematic of the four-stage pipeline underlying the V1 and V2 notebooks: (i) environment setup, including repository installation and CUDA validation; (ii) interactive parameter generation, converting structured user input into a Boltz-2-compliant YAML configuration; (iii) the prediction engine, which invokes the Boltz-2 command-line tool; and (iv) automated visualization and analysis of structural and affinity outputs. Arrows indicate file-based data flow between stages (params.yaml, run_params.txt, and job-specific output directories); each stage can be inspected or re-executed independently within the same notebook session.

Data flow between stages is file-based rather than held in memory across cells, which allows the pipeline to be resumed or partially re-executed without repeating earlier steps. The parameter-generation stage writes two artifacts to the working directory - a Boltz-2-compliant params.yaml and a companion run_params.txt recording execution settings - which are read by the prediction stage; the prediction stage in turn writes structure files (PDB/CIF), confidence JSON, and pLDDT/PAE arrays to a job-specific output directory, which the analysis stage parses without needing to know how the underlying prediction was produced. This decoupling means the analysis and visualization code is agnostic to whether a job originated from the single-job or batch pipeline, and it is what allows the batch notebook (§2.6) to reuse the same downstream analysis logic while replacing only the orchestration layer above it.

It is important to distinguish, throughout this section, between functionality that Boltz2-Notebook implements and functionality it merely exposes: the underlying structure and affinity-prediction model, its YAML input schema, and its confidence/affinity output format are all inherited unmodified from the boltz package (Passaro et al., 2025; Wohlwend et al., 2024). Boltz2-Notebook’s contribution is the software layer around that model - input construction, execution management, and output interpretation - not any change to the model itself.

### 2.2 Environment Setup

The setup stage (setup.py) replaces the manual installation procedure normally required to run Boltz-2 - creating a Python environment, installing a CUDA-matched PyTorch build (Paszke et al., 2019), cloning and installing the boltz package with its optional CUDA extras, and installing auxiliary dependencies (Biopython (Cock et al., 2009), NumPy (Harris et al., 2020), Matplotlib (Hunter, 2007), PyYAML, py3Dmol) - with a single automated cell. Execution proceeds through three scripted steps: repository cloning, dependency installation via pip install -e boltz[cuda] together with the auxiliary packages, and a validation step that imports PyTorch and reports CUDA availability and device count. Each step is wrapped in explicit success/failure handling so installation failures are reported with the responsible step identified rather than surfacing as an unstructured traceback, and any pre-existing installation directory from a prior session is removed before cloning to avoid stale-state failures on notebook re-runs. Since this stage runs inside the Google Colab runtime (Bisong, 2019), it runs on a GPU-backed virtual machine provisioned by Colab (typically an NVIDIA T4) without requiring the user to own or configure GPU hardware, install CUDA drivers, or manage a local Python environment - the three most common points of failure in a manual Boltz-2 installation.

### 2.3 Interactive Parameter Generation

The most direct attempt to address the Boltz-2 usability barrier within Boltz2-Notebook is the parameter-generation stage (param_gen.py), which is also the clearest software-engineering contribution of this work. Boltz-2’s native input format is a YAML document with a specific, nested schema. It consists of: 1) a sequences block containing a list of typed entities (protein, DNA, RNA, ligand), 2) an optional constraints block encoding covalent bonds, pocket definitions, and contact restraints, 3) an optional templates block referencing external structure files, and 4) a properties block specifying the chain for which to predict the affinity. This schema is expressive, but strict - a single malformed indentation, a mistyped field, or an invalid residue/atom reference will cause the prediction to fail, often with an uninformative error message. Building it by hand requires familiarity with both YAML syntax and Boltz-2’s internal conventions.

param_gen.py replaces hand-authored YAML files with a structured, form-driven build process. Each biomolecular entity - protein, DNA or RNA sequence or small-molecule ligand provided as either a SMILES string (Weininger, 1988) or a PDB Chemical Component Dictionary (CCD) code (Westbrook et al., 2015) - is entered by users in individual input fields. Dedicated helper functions normalise sequences, sanitise chain identifiers and validate the entity before it is added to the internal data structure. Post-translational modifications are specified as (position, CCD code) pairs attached to a given protein chain, cyclic chains are specified by a boolean rather than requiring the user to encode chain topology manually, and a custom multiple sequence alignment (MSA) file can be supplied per protein chain instead of server-generated alignment. Structural constraints such as covalent bonds between named atoms, pocket definitions specifying binder and contact-residue sets with a distance threshold, and pairwise contact restraints with soft or hard enforcement are built using the same structured interface instead of requiring the user to hand-write the corresponding YAML blocks (Table 2). Template-guided modelling is enabled by allowing users to upload a reference PDB or CIF structure, select the chains to which this applies, and specify a confidence threshold for how strongly the template constrains the prediction.

The assembled configuration is internally serialised to params.yaml with a custom YAML dumper that has explicit representers for identifier lists and quoted scalars, such that chain identifiers, SMILES strings, and sequences are emitted in a form accepted by Boltz-2’s parser without post-processing. Execution settings (recycling steps, sampling steps, diffusion samples, MSA parameters) are separately recorded in run_params.txt and consumed by the prediction stage. This separation of biomolecular input from execution configuration allows the same input specification to be re-run with different sampling or refinement settings without the need to reconstruct the entities and constraints from scratch. The practical advantage of this design over manual editing of YAML is that the syntactic and structural validity of the generated configuration are guarantyd by construction: the user only interacts with domain level fields (a sequence, a residue position, a distance threshold) and the schema-level correctness of the resulting file is the responsibility of the software, not the user.

### 2.4 Prediction Engine

The execution stage (Boltz_Run.py) reads the run-configuration file written in §2.3 and calls the boltz predict command-line tool with the appropriate arguments, including the number of recycling steps, diffusion sampling steps, diffusion samples, MSA pairing strategy and maximum MSA depth, using the MSA server hosted by Boltz-2 unless a custom alignment was provided. Execution output is piped through a lightweight logging layer that strips out ANSI terminal control codes and displays progress using an animated status indicator, so that long-running predictions provide continuous, human-readable feedback within the notebook, instead of raw CLI output. Job-level parameters—recycling steps, sampling steps, diffusion samples, step scale, MSA depth, and an optional physics-based potentials refinement—are exposed as user-configurable settings with documented defaults, such that a single execution stage can serve three broad use profiles (fast, balanced and high-quality; Table 3) by adjusting sampling depth and refinement rather than requiring separate code paths. Each job writes its outputs to a dedicated subdirectory named after the job, containing predicted structures, per-model confidence and affinity JSON files, and pLDDT/PAE arrays. An override flag controls whether to reuse an existing output directory (skipping re-computation) or replace it. Model execution failures are caught at the subprocess level and reported with the corresponding job identifier, rather than terminating the notebook session. This is particularly important for the batch pipeline described in §2.6, where a single failed target should not stop processing of subsequent targets.

### 2.5 Visualization and Analysis

Analysis stage (analysis.py) takes the structure and confidence outputs from §2.4 and converts them into a combined set of visual and tabular summaries. The user does not need to write any post-processing code. Per-residue confidence (pLDDT) and predicted aligned error (PAE) are extracted from the native output arrays of Boltz-2 and visualised as colour-coded overlays on an interactive three-dimensional structure viewer (via py3Dmol), allowing confidence to be inspected in structural context rather than as an isolated numeric summary (Rego & Koes, 2015). For jobs with an affinity-designated binder chain, the reported affinity value and binding-probability score of Boltz-2 are extracted and passed through classification functions that map the continuous scores onto qualitative categories according to threshold conventions documented in the software’s user-facing guide. These are compiled into a single dashboard panel that integrates a binding-probability summary, affinity-strength summary and structure-quality indicators for a given job. All confidence and affinity outputs and generated plots are further organised into a per-job HTML report and a structured summary object, which can be exported or aggregated downstream. The underlying confidence and affinity metrics are themselves entirely computed by Boltz-2, like the constraint schema in §2.3; this stage’s role is to extract, visualise and organise them into an interpretable per-job summary, rather than raw output files scattered across a directory tree.

### 2.6 Batch Processing

The batch notebook refactors the single-job pipeline into a seven-stage, manifest-driven workflow (Figure 2) designed for screening multiple targets in a single execution session, a use case not supported by the base boltz CLI without external scripting. Input can be supplied in CSV or FASTA formats. The target is a list of targets, or pre-written YAML files provided individually or in a compressed archive, and is transformed into a validated job manifest before any prediction. This preflight validation step ensures the manifest is complete and flags malformed entries before computationally expensive jobs are launched, rather than letting failures appear only after the execution starts. Jobs are executed on a resumable queue that tracks the status of each target, so that an interrupted batch run can be resumed without recomputing targets that have already finished. It also supports configurable run profiles that balance the depth of prediction against the throughput of the batch. When completed, results for all targets are merged and ranked according to user-selected metrics (affinity probability, confidence score, pLDDT, or predicted template modelling score), and packaged into a single exportable archive along with job logs, facilitating easy triage of screening results and post hoc inspection of failed or low-confidence targets.

**Figure 2.**
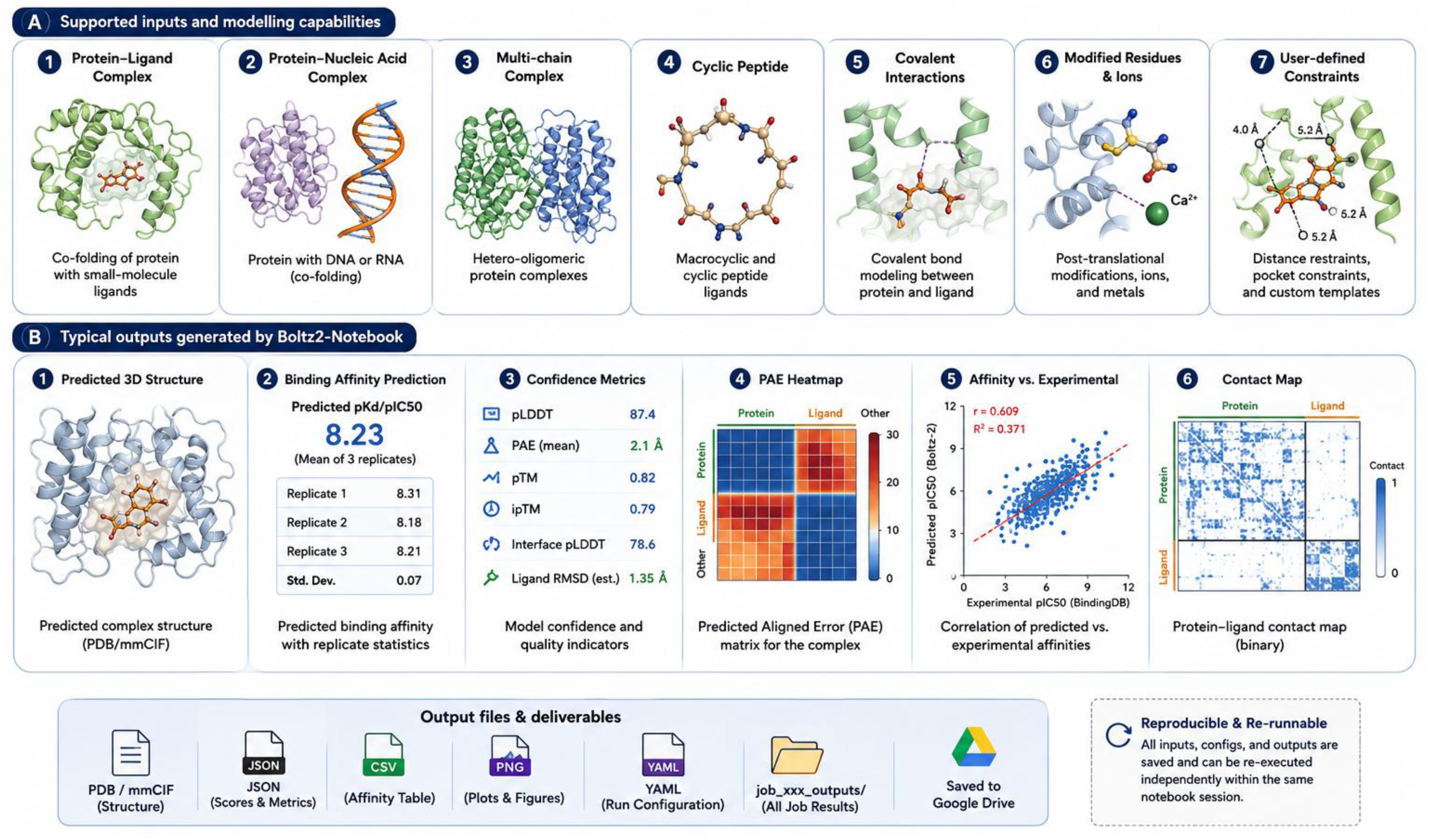
Batch processing pipeline. Seven-stage workflow implemented by the batch notebook: input handling (CSV/FASTA/YAML manifest construction), preflight validation, launch/resume of queued execution, per-target result summarization and ranking, structure visualization, job-log/failure diagnostics, and single-archive export. The batch pipeline reuses the parameter-generation and analysis logic of the single-job notebooks (Figure 1) while replacing single-job orchestration with resumable, multi-target queue management.

### 2.7 Software Availability

The Boltz2-Notebook is an open-source code released under the MIT license and is available free of charge on GitHub (github.com/AtharvaTilewale/boltz2-notebook) with a project website documenting the available notebooks, comparison of features between different versions and usage instructions. All three notebooks (V1, V2 and Batch) can be run directly from the repository using Google Colab without local installation, in line with the accessibility goal discussed in the Introduction. The repository contains source code for all four pipeline stages, allowing the parameter-generation, execution, and analysis logic described above to be inspected, modified, or reused independently of the notebook interface. Supplementary Table S4 provides full software metadata (version, dependencies, license and support information).

## 3. Software Features and Applications

### 3.1 Accessibility and Ease of Use

The main practical difference between Boltz2-Notebook and the base boltz package is not modelling capability but rather the effort that is needed to achieve a first prediction. Execution is performed only within the Google Colab runtime, so the only requirement for a user is a web browser and a Google account, with no need for a local NVIDIA GPU, CUDA toolkit, or Python environment and the steps of dependency installation, version compatibility, and matching with CUDA that normally precede any Boltz-2 run (§2.2) are automatically performed at the beginning of the session. The interactive notebook format also has the effect that input specification, execution and result interpretation takes place in a single, linear sequence of cells, rather than as separate command line invocations, with manually managed intermediate files. For a user without a background in computational biology or software engineering, e.g. an experimental structural biologist or a student encountering diffusion-based structure prediction for the first time, this lowers the practical entry cost from setting up a GPU-enabled scientific computing environment to opening a shared link and providing a sequence or SMILES string. This is an accessibility improvement in the same sense as ColabFold for AlphaFold2 (Mirdita et al., 2022): the underlying predictive ability is unchanged, but the population of researchers who can exercise it without local infrastructure or command-line proficiency is substantially larger.

### 3.2 Comprehensive Biomolecular Modeling

Boltz2-Notebook exposes the full range of biomolecular entity types supported by Boltz-2: single-chain and multi-chain proteins, protein-ligand complexes with ligands specified as SMILES strings or CCD codes, and, in the V2 notebook, DNA and RNA sequences alongside proteins within the same complex. All of these modeling capabilities - the underlying diffusion architecture, the co-folding of heterogeneous entity types, and the affinity-prediction head for designated binder chains - originate entirely from the Boltz-2 model itself (Passaro et al., 2025) and are unmodified by this work. What Boltz2-Notebook provides is a uniform interface for specifying and combining these entities: a user assembling a multi-chain antibody-antigen complex, a nucleic-acid-bound transcription factor, or a simple protein-ligand pair interacts with the same structured input fields regardless of entity type or complex composition, rather than needing to hand-construct correspondingly different YAML blocks for each case. This is a usability distinction, not an expansion of what can be modeled.

### 3.3 Advanced Prediction Configuration

Beyond basic complex assembly, Boltz-2’s schema supports a set of structure-guiding mechanisms that are of particular relevance to structure-based drug design and comparative modeling: template-guided prediction, in which a reference structure biases the output toward a known fold or binding mode; pocket constraints, which restrict a ligand or binder to a defined set of contact residues within a distance threshold; contact restraints, which enforce or encourage proximity between arbitrary residue or atom pairs; covalent-bond constraints, for modeling disulfides, covalent inhibitors, or engineered linkages; residue modifications, specified as CCD-coded post-translational modifications at defined sequence positions; cyclic-chain topology, for head-to-tail cyclized peptides; and multiple sequence alignment control, allowing either automatic server-based MSA generation or the substitution of a user-supplied, pre-computed alignment. Each of these mechanisms is a native Boltz-2 capability, described in the model’s input specification; Boltz2-Notebook’s contribution, as detailed in §2.3, is exposing them through discrete, validated input fields - a residue index and CCD code for a modification, a distance value and residue list for a pocket - rather than requiring the user to construct the corresponding nested YAML syntax by hand. Because misconfiguration of these constraint blocks is a common source of silent failure or unintended model behavior in manually authored inputs, the practical effect of this design is to make advanced, constraint-guided prediction accessible to users who would otherwise be limited to unconstrained, default-configuration runs.

### 3.4 Automated Analysis and Visualization

Reading a structure prediction is more than just the coordinate file itself: per-residue confidence, inter-residue error estimates, and if applicable, affinity and binding-probability scores, all feed into the question of whether a given model is trustworthy enough to act on. Boltz2-Notebook automates extraction and presentation of these quantities immediately after prediction. Predicted structures are visualised in an interactive 3-D viewer with pLDDT mapped onto a per-residue colour scale, making it possible to evaluate confidence in structural context, e.g. telling apart a well-resolved binding pocket from a low-confidence flexible loop in the same model, rather than as a single averaged number. The predicted aligned error is displayed as a two-dimensional matrix, allowing for examination of inter-domain or inter-chain positional confidence relevant for multi-chain assemblies. If a binder chain has been specified for affinity prediction, the affinity value and binding-probability score are reported along with these structural confidence metrics in a single dashboard view. All figures generated during analysis are saved to the job output directory in a format suitable for direct inclusion in a manuscript or report. As in §3.3, Boltz-2 itself computes all the fundamental metrics - pLDDT, PAE, affinity value, binding probability. The notebook contribution is their aggregation into an interpretable, per-job visual summary, lowering the analysis burden from writing custom plotting and parsing code against Boltz-2’s raw JSON/NPZ outputs to reviewing an automatically generated report.

### 3.5 Reproducibility and Workflow Management

Each prediction job is written to a dedicated, consistently structured output directory containing the generated input configuration (params.yaml, run_params.txt), predicted structures, confidence and affinity outputs, and derived figures, so that the parameters used to produce a given result remain attached to that result rather than existing only in transient notebook state. This organization allows a completed job to be re-inspected or re-derived without re-running the prediction, and allows a user to confirm precisely which sampling settings, constraints, and MSA source produced a given structure. Optional integration with Google Drive allows this output structure to be persisted beyond the lifetime of the Colab session, and results can additionally be exported as a single compressed archive containing all job outputs and configuration files. These properties are relevant to reproducibility in the practical sense emphasized in current bioinformatics software guidance: a collaborator or reviewer supplied with a job’s exported archive has direct access to the exact configuration that generated it, rather than needing to reconstruct the run from a text description of parameters used.

### 3.6 High-Throughput Batch Prediction

Where the single-job notebooks are appropriate for investigating individual targets or complexes, the batch notebook (§2.6) is designed for screening or dataset-scale use cases in which the same prediction workflow must be applied to many targets - for example, evaluating a candidate compound library against a fixed protein target, or generating structures and affinity estimates across a curated set of protein-ligand pairs, as in the benchmark described later in this manuscript. By accepting CSV, FASTA, or bundled YAML input, validating the full job set before execution begins, and supporting resumable, queued processing of large target lists, the batch pipeline removes the need for external scripting to iterate the single-job workflow across many inputs – a task that would otherwise require the user to independently manage job submission, failure handling, and result aggregation using the base boltz CLI. The resulting ranked, aggregated output is intended to support triage-style analysis, in which a large candidate set is narrowed to a smaller number of high-confidence or high-affinity targets for further, more detailed inspection using the single-job notebooks.

### 3.7 Example Applications

Within the scope of what Boltz-2 itself supports, Boltz2-Notebook is applicable to a range of practical structural biology tasks: predicting protein-ligand complex structures and estimating relative binding affinity for candidate compounds in early-stage drug discovery; screening modest-sized compound sets against a target of interest using the batch workflow prior to more computationally intensive downstream evaluation; generating structural hypotheses for engineered or mutated protein variants, optionally guided by a related template structure; modeling protein-DNA/RNA or multi-chain protein assemblies for mechanistic hypothesis generation; and, given its low barrier to entry, use in graduate-level teaching or workshop settings where participants require hands-on exposure to modern structure-prediction methods without institutional GPU access. These applications reflect the intended use cases for which the underlying accessibility and configuration features described above were designed, rather than capabilities specific to or validated exclusively within this work; the extent to which affinity predictions generated through this or any Boltz-2-based workflow can be relied upon for prioritization decisions is addressed directly in the independent benchmark presented later in this manuscript.

## 4. Benchmark Dataset and Methods

### 4.1 Benchmark Design

Boltz-2’s original description (Passaro et al., 2025) reports strong binding-affinity prediction performance on internally curated benchmarks, including the FEP+ hit-to-lead/lead-optimization benchmark, the CASP16 affinity assessment track, and the MF-PCBA hit-discovery benchmark. Independent evaluation on data assembled separately from a model’s own training and benchmarking pipeline is a necessary complement to developer-reported results, both because it probes generalization to differently sourced and differently curated data, and because it permits assessment of properties - such as the reliability of self-reported confidence metrics as indicators of prediction accuracy - that are not always characterized in the original publication. We therefore designed and executed an independent benchmark of Boltz-2’s protein-ligand binding-affinity prediction performance using a newly curated dataset drawn from BindingDB (Liu et al., 2007), a large, continuously updated, literature-curated repository of experimentally measured protein-ligand binding data.

This benchmark is presented as a methodologically distinct component of the present work. It evaluates the predictive performance of the underlying Boltz-2 model as released, and is not a validation of the Boltz2-Notebook software described in Sections 2 and 3. Prediction jobs for the benchmark were executed directly through the Boltz-2 command-line interface on a high-performance computing cluster, using the same YAML input schema and prediction engine that Boltz2-Notebook automates and exposes through its interactive interface, but without routing execution through the notebook itself (Section 4.6). The benchmark therefore demonstrates the input specification and prediction workflow that Boltz2-Notebook automates for its users, while the reported performance metrics reflect the accuracy of Boltz-2 itself rather than any property specific to the notebook implementation.

### 4.2 Dataset Collection

The benchmark dataset was derived from BindingDB’s curated ‘Ligand-Target Affinity’ bulk release (Gilson et al., 2016; Liu et al., 2025; Liu et al., 2007), comprising 94,220 entries at the time of retrieval. This initial dataset was reduced to a final benchmark set through a two-stage, script-based curation pipeline designed to enforce structural, chemical, and experimental consistency (Figure 3).

**Figure 3.**
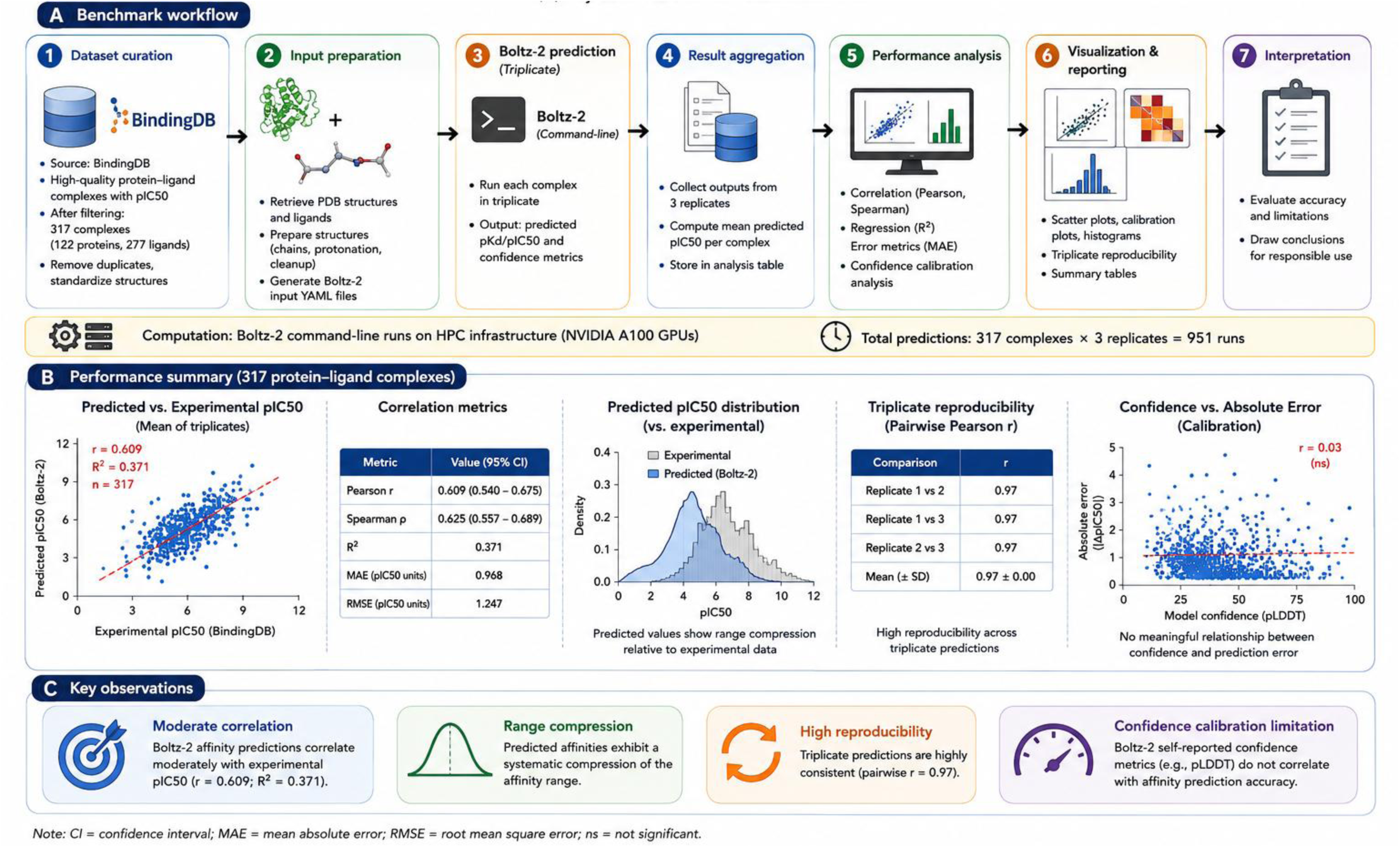
Curation pipeline for the independent BindingDB benchmark dataset. Flow diagram illustrating stepwise reduction of the initial BindingDB ‘Ligand-Target Affinity’ bulk release (94,220 entries) to the final benchmark dataset (317 protein-ligand pairs), showing the number of entries retained after each filtering step: single-chain/single-PDB target restriction, mandatory-field completeness, valid quantitative affinity measurement, valid numeric IC50, RDKit-parseable ligand structure (≤128 heavy atoms), protein sequence validity (≤1,800 residues), duplicate removal, and the per-protein ligand cap (≤3 ligands per target).

In the first stage, entries were retained only if the associated target was annotated as a single-chain protein with exactly one associated PDB identifier (Berman et al., 2000); entries with multiple, comma-, semicolon-, or pipe-separated PDB identifiers, or with more than one annotated protein chain, were excluded to avoid ambiguity in structural representation. Entries missing any of five mandatory fields - BindingDB identifier, ligand SMILES, target protein name, PDB identifier, and protein sequence - were removed. Entries were further required to report at least one quantitative affinity measurement among IC50, EC50, or Kd.

In the second stage, entries were restricted to those reporting a valid, strictly positive numeric IC50 value; entries with missing, non-numeric, malformed, or non-positive IC50 values were discarded. Ligand structures were validated using RDKit (Landrum, 2006), retaining only entries with a chemically parseable SMILES string corresponding to a molecule of no more than 128 heavy atoms. Protein sequences were validated against the standard amino-acid alphabet (including ambiguity codes) and restricted to a maximum length of 1,800 residues. Exact-duplicate records were removed, and, to limit the influence of individual over-represented targets on aggregate statistics, a maximum of three ligand entries was retained per protein, with duplicate (protein, ligand) pairs for a given target removed prior to this cap being applied.

This pipeline yielded a final benchmark dataset of 317 protein-ligand pairs, comprising 122 unique protein sequences and 277 unique ligand structures, each associated with a single experimentally measured IC50 value and a single corresponding PDB structure identifier. The complete curation pipeline is implemented as two standalone, reusable Python scripts (curation and cleaning), provided in the Supplementary Material together with the resulting dataset, to allow independent reproduction of the filtering procedure from the original BindingDB bulk release.

### 4.3 Prediction Workflow

Each of the 317 curated entries was converted into a Boltz-2-compliant YAML input file, following the schema described in Section 2.3, comprising a protein entity (chain A, defined by the curated sequence) and a ligand entity (chain B, defined by the curated SMILES string), with the ligand chain designated as the binder for affinity prediction. YAML generation for the benchmark was performed by a dedicated conversion script that applies the same sequence-cleaning and quoting conventions used by param_gen.py (Section 2.3), producing one input file per BindingDB entry together with a manifest recording the source identifier, protein length, and ligand SMILES for each generated job.

Prediction execution was performed using the boltz predict command-line interface directly, rather than through the Colab-based notebook pipeline described in Section 2.4, in order to allow batched, unattended execution of all 317 jobs on dedicated GPU compute (Section 4.6). Multiple sequence alignments were generated automatically via the Boltz-2 MSA server for each target (--use_msa_server); no custom or pre-computed alignments were supplied. Structure sampling was performed with 250 diffusion sampling steps and a single structural sample per job, consistent with standard-quality single-structure prediction; affinity prediction was performed with 250 affinity sampling steps and 10 affinity diffusion samples per job, following Boltz-2’s recommended configuration for affinity estimation. Each of the 317 targets was predicted in triplicate - three independent prediction runs per protein-ligand pair, differing only in the model’s internal stochastic sampling - to allow assessment of run-to-run prediction stability (Section 4.5) independently of dataset-level accuracy.

Following execution, per-job confidence JSON files, affinity JSON files, and per-residue pLDDT, PAE, and predicted distance error (PDE) arrays were programmatically collected and consolidated into a single tabular dataset using a dedicated aggregation script. Boltz-2 reports affinity as a continuous, log-scale predicted value; this value was converted, for each replicate, to a predicted IC50 (μM), a predicted pIC50, and a predicted binding free energy (kcal mol⁻¹) using the fixed transformation documented for Boltz-2’s affinity output, and averaged across the three replicates per target to obtain a single consensus predicted value for each protein-ligand pair, alongside the corresponding replicate standard deviation and coefficient of variation. Experimentally reported IC50 values were converted to pIC50 on the same scale to allow direct comparison with predicted values. The complete set of per-replicate and consensus values, together with all associated confidence and structural-quality metrics, constitutes the analyzed benchmark dataset used for the statistical evaluation described below (Supplementary Table S1); per-replicate values prior to consensus averaging are additionally reported in Supplementary Table S2.

### 4.4 Statistical Evaluation

Predictive performance was evaluated by comparing the consensus (replicate-averaged) predicted pIC50 against the experimentally reported pIC50 for all 317 benchmark entries, using the following metrics.

Pearson correlation coefficient (r), quantifying the linear association between predicted and experimental values, was computed in the standard form 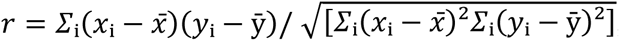, where x_i_ and y_i_ denote the experimental and predicted pIC50 values, respectively, for entry i.

Spearman rank correlation coefficient (ρ), quantifying the monotonic association between predicted and experimental values based on their rank order, was computed as the Pearson correlation of the ranked variables.

Coefficient of determination (R^2^) was reported as the square of the Pearson correlation coefficient for the linear relationship between predicted and experimental pIC50.

Mean absolute error (MAE) was computed as *MAE* = (1/*n*)*Σ*_i_|ŷ_i_ − *y*_i_|, and root mean squared error (RMSE) as 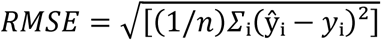, where ŷ_i_ and y_i_ denote the predicted and experimental pIC50 values for entry i, and n = 317.

To characterize the precision of these point estimates, bootstrap confidence intervals were computed following the nonparametric resampling procedure of Efron (Efron, 1979), by resampling the 317 protein-ligand pairs with replacement, recomputing each of the above statistics on each resampled dataset, and reporting the 2.5th and 97.5th percentiles of the resulting distribution as an approximate 95% confidence interval.

Residual analysis was performed on the signed prediction error (predicted minus experimental pIC50) as a function of experimental pIC50, to assess systematic bias, including potential range compression at the extremes of the affinity scale, and heteroscedasticity in prediction error across the affinity range. A complementary Bland-Altman analysis (Bland & Altman, 1986) was performed by plotting the mean of the predicted and experimental pIC50 for each entry against their signed difference, with the mean bias and 95% limits of agreement (bias ± 1.96 × SD of the differences) reported to characterize systematic and random components of disagreement between predicted and experimental values.

Calibration analysis was performed to assess whether Boltz-2’s self-reported confidence metrics (confidence score, interface predicted TM-score, complex pLDDT) are informative of prediction accuracy. Entries were grouped into bins by each confidence metric, and the mean absolute prediction error was computed within each bin; the association between binned confidence and mean absolute error was quantified using the Pearson correlation between the continuous confidence metric and the per-entry absolute error, providing a reliability-diagram-style assessment analogous to calibration evaluation in classification settings.

### 4.5 Benchmark Quality Control

Prediction stability was assessed using the triplicate design described in Section 4.3. For each protein-ligand pair, the coefficient of variation of the three replicate predicted IC50 values was computed, and pairwise Pearson correlations between replicates (replicate 1 vs. 2, 1 vs. 3, and 2 vs. 3) were calculated across the full dataset to characterize overall run-to-run reproducibility of Boltz-2’s stochastic sampling procedure independent of agreement with experimental data.

Job-level execution failures were handled at the level of the batch submission script (Section 4.6): each target was submitted as an independent job, and failure of an individual job did not halt processing of subsequent targets in the batch. Job status (success, failure, or skipped owing to pre-existing output) was recorded for every submitted target in a structured status log. Targets for which no valid confidence or affinity output was produced were excluded from the final aggregated dataset rather than imputed, and the aggregation script used to construct the analyzed dataset (Section 4.3) discovers and validates the presence of the expected confidence JSON, affinity JSON, and structural output files for each job prior to inclusion, so that the final 317-entry dataset consists exclusively of targets with complete, successfully generated prediction output across all three replicates.

### 4.6 Computational Environment

Benchmark predictions were executed on the University of Edinburgh’s Eddie high-performance computing cluster, using a single GPU-backed compute node per job (-l gpu=1) with CUDA 12.1.1, rather than within the Google Colab environment used for the interactive Boltz2-Notebook workflow described in Sections 2 and 3; this distinction is noted explicitly because it separates the computational environment used to generate the benchmark results reported in this manuscript from the environment in which Boltz2-Notebook itself is intended to be used by end users. Job submission and resource allocation were managed through the cluster’s Grid Engine scheduler, with per-job wall-clock time, memory, and GPU allocation specified in the associated submission script (Supplementary Material). The Boltz-2 prediction engine and its Python dependencies were installed in a dedicated virtual environment on the cluster’s scratch filesystem, with a persistent model-weight cache shared across all 951 (317 × 3) prediction jobs to avoid redundant downloads. The specific boltz package version and complete dependency manifest used for the benchmark are reported in the Supplementary Material to support exact reproduction of the reported results; all data curation, YAML generation, and result-aggregation scripts described above are provided alongside the benchmark dataset in the associated code repository.

## 5. Results

### 5.1 Benchmark Dataset

We applied the two-stage curation pipeline (Section 4.2) to the initial BindingDB pull of 94,220 entries, resulting in a final benchmark dataset of 317 protein-ligand pairs (Figure 3) with 122 unique protein sequences and 277 unique ligand structures, each with a single experimentally measured IC50 value and a single corresponding PDB structure identifier. Protein sequence lengths ranged from 147 to 1,560 residues (mean 588, median 491) and the number of ligand heavy atoms (as computed by RDKit) ranged from 5 to 122 atoms (mean 27.4, median 26), consistent with the ≤1,800-residue and ≤128-atom filtering criteria used during curation. Experimental potency ranged across 8 orders of magnitude from pIC50 1.74 to 9.64 (approximately 18 nM to 180 mM in linear IC50 terms). Table 4 shows summary statistics of the dataset.

### 5.2 Overall Affinity Prediction Performance

Comparing the replicate-averaged predicted pIC50 against the experimental pIC50 across all 317 benchmark entries (Figure 4) yielded a Pearson correlation coefficient of r = 0.609 (95% bootstrap CI: 0.540-0.675) and a Spearman rank correlation of ρ = 0.625 (95% CI: 0.552-0.691). The coefficient of determination was R^2^ = 0.371 (95% CI: 0.291-0.455). Prediction error was MAE = 0.968 pIC50 units (95% CI: 0.882-1.054) and RMSE = 1.239 pIC50 units (95% CI: 1.135-1.345). All confidence intervals were computed from 5,000 bootstrap resamples of the 317 protein-ligand pairs. Summary statistics with associated confidence intervals are reported in Table 5.

**Figure 4.**
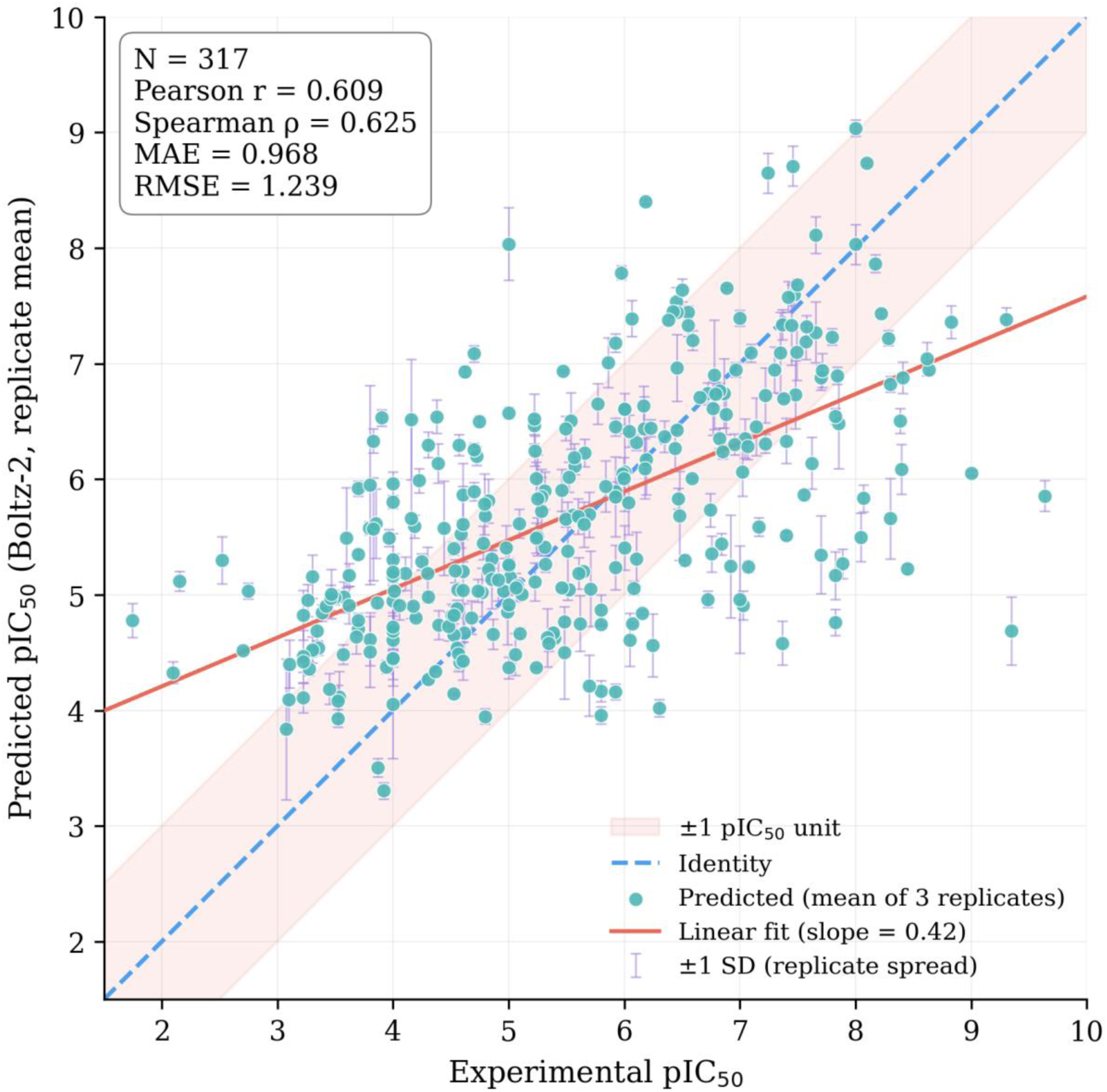
Predicted versus experimental binding affinity across the benchmark dataset. Scatter plot of replicate-averaged Boltz-2-predicted pIC50 against experimentally reported pIC50 for all 317 benchmark entries. The dashed line indicates identity (perfect agreement); the solid line indicates the linear regression fit; the shaded band indicates ±1 pIC50 unit from identity. Inset reports Pearson correlation coefficient (r), Spearman correlation coefficient (ρ), mean absolute error (MAE), and root-mean-squared error (RMSE); 95% bootstrap confidence intervals for all metrics are reported in Table 5.

Predicted pIC50 values occupied a narrower range (3.31-9.04) than experimental values (1.74-9.64). Of the 317 entries, 104 (32.8%) had an absolute prediction error below 0.5 pIC50 units (approximately 3-fold in linear IC50), 189 (59.6%) had an absolute error below 1.0 pIC50 unit (approximately 10-fold), and 31 (9.8%) had an absolute error exceeding 2.0 pIC50 units.

### 5.3 Reproducibility and Prediction Stability

Across the triplicate predictions generated for each of the 317 targets, pairwise Pearson correlation between replicate predicted pIC50 values was r = 0.97 for all three replicate pairs (replicate 1 vs. 2, 1 vs. 3, and 2 vs. 3; Figure 5). The coefficient of variation of the three replicate predicted IC50 values per target had a mean of 31.4% and a median of 25.7% across the dataset, with values ranging from 0.5% to 157.3% (full per-target values in Supplementary Table S3). No significant association was observed between replicate coefficient of variation and absolute prediction error against experimental data (r = 0.057, p = 0.31; mean absolute error by CV quartile: 0.94, 1.02, 0.93, 0.98 pIC50 units, from lowest to highest CV quartile).

**Figure 5.**
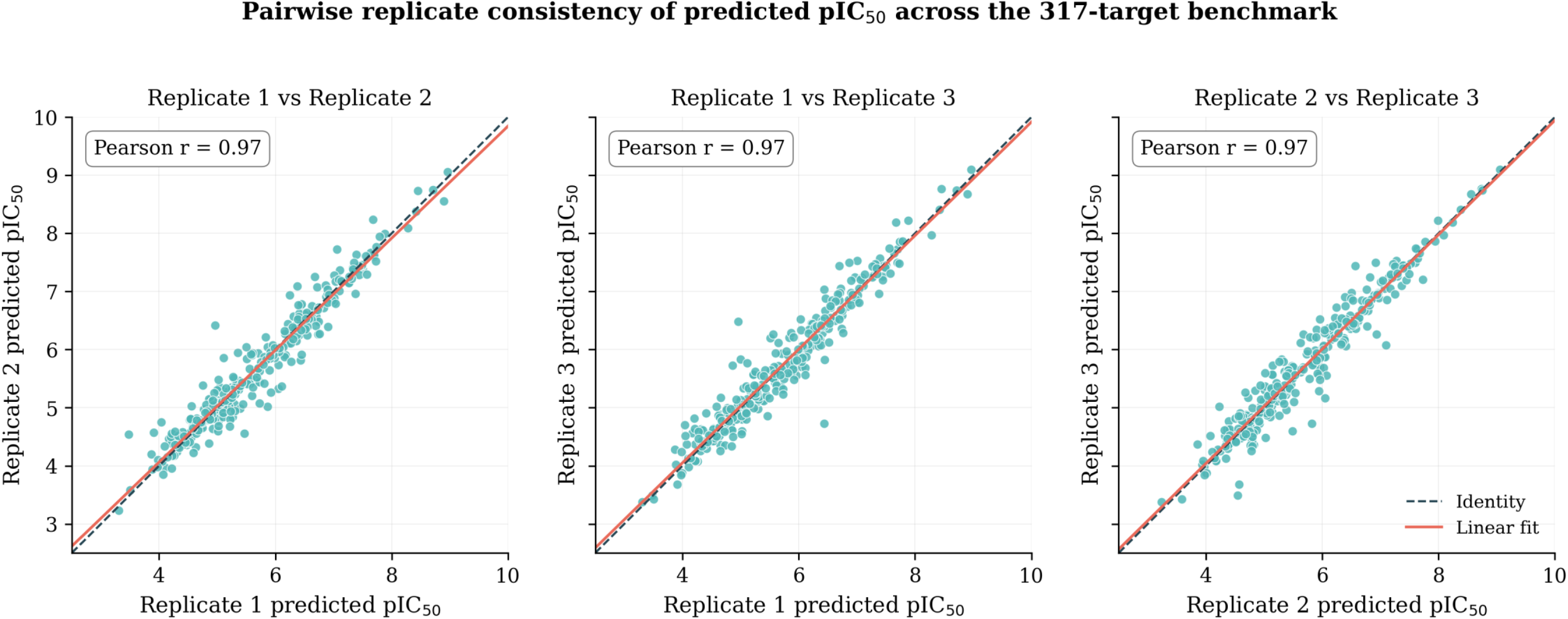
Triplicate replicate consistency of predicted binding affinity. Pairwise scatter plots comparing replicate-predicted pIC50 values across the three independent prediction runs performed for each of the 317 benchmark targets: replicate 1 vs. replicate 2, replicate 1 vs. replicate 3, and replicate 2 vs. replicate 3 (left to right). The dashed line indicates identity; the solid line indicates the linear regression fit. Pearson correlation coefficients for each replicate pair are reported inset.

No target failed to produce complete output across all three replicates following the quality-control filtering described in Section 4.5; the final 317-entry dataset therefore reflects targets with fully successful triplicate prediction.

### 5.4 Confidence Assessment

Boltz-2’s self-reported structural and interface confidence metrics were, on average, high across the benchmark: mean confidence score 0.814 (SD 0.121, range 0.381-0.976), mean interface predicted TM-score (iPTM) 0.882 (SD 0.125, range 0.363-0.994), and mean complex pLDDT 0.797 (SD 0.140, range 0.370-0.981).

No significant linear relationship was observed between any of these confidence metrics and absolute affinity prediction error: confidence score (r = 0.040, p = 0.48), iPTM (r = −0.007, p = 0.91), and complex pLDDT (r = 0.045, p = 0.43). Binning entries into quartiles by confidence score, mean absolute error was 1.03, 0.76, 1.02, and 1.07 pIC50 units from the lowest to the highest confidence quartile (Figure 6); binning by complex pLDDT quartile, mean absolute error was 0.94, 0.87, 0.93, and 1.13 pIC50 units from lowest to highest pLDDT quartile. In both cases, the highest-confidence quartile did not show the lowest mean absolute error.

**Figure 6.**
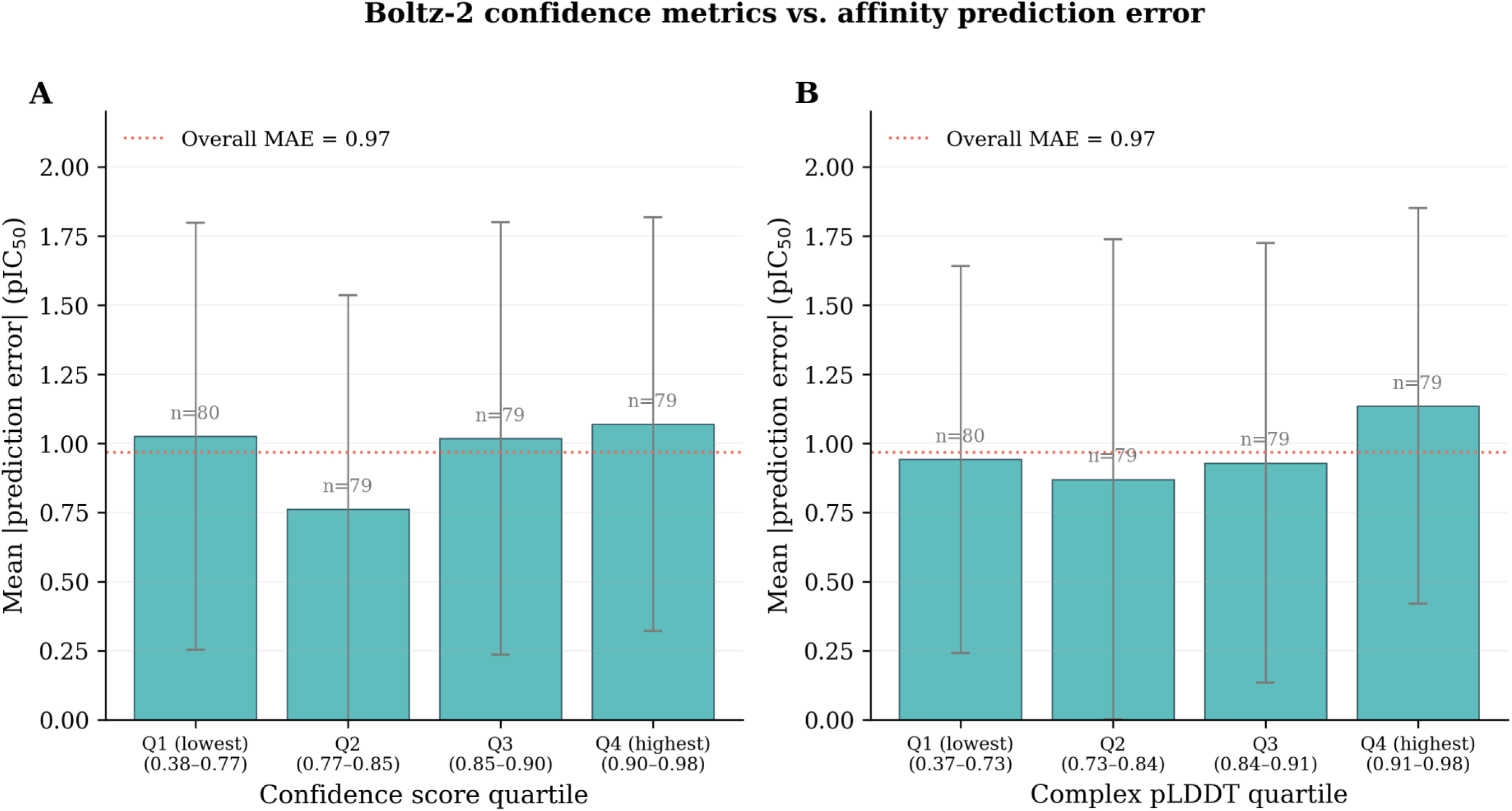
Relationship between Boltz-2 confidence metrics and affinity prediction error. Reliability-diagram-style summary of mean absolute affinity prediction error (pIC50 units) binned by quartile of Boltz-2’s self-reported confidence score (A) and complex pLDDT (B), across the 317-entry benchmark. Error bars indicate the standard deviation of absolute error within each quartile bin. The absence of a monotonic decrease in error from the lowest to the highest confidence/pLDDT quartile indicates that these metrics do not linearly track affinity prediction accuracy in this benchmark (Pearson correlation between each metric and absolute error reported in the main text).

### 5.5 Error Analysis

Signed prediction error (predicted minus experimental pIC50) was significantly correlated with experimental pIC50 (r = −0.725, p = 5.0 × 10⁻⁵³), indicating a systematic relationship between the direction and magnitude of error and the experimental potency of the entry: high-potency compounds (high experimental pIC50) tended to be underpredicted, and low-potency compounds tended to be overpredicted (Figure 7A). Splitting the dataset at the median experimental pIC50, the Pearson correlation within the lower-potency half (n = 158) was r = 0.349, and within the higher-potency half (n = 159) was r = 0.349 - both substantially lower than the pooled correlation (r = 0.609) computed across the full dynamic range. Mean absolute error was comparable between the two halves (0.982 vs. 0.954 pIC50 units for the lower-and higher-potency halves, respectively).

**Figure 7.**
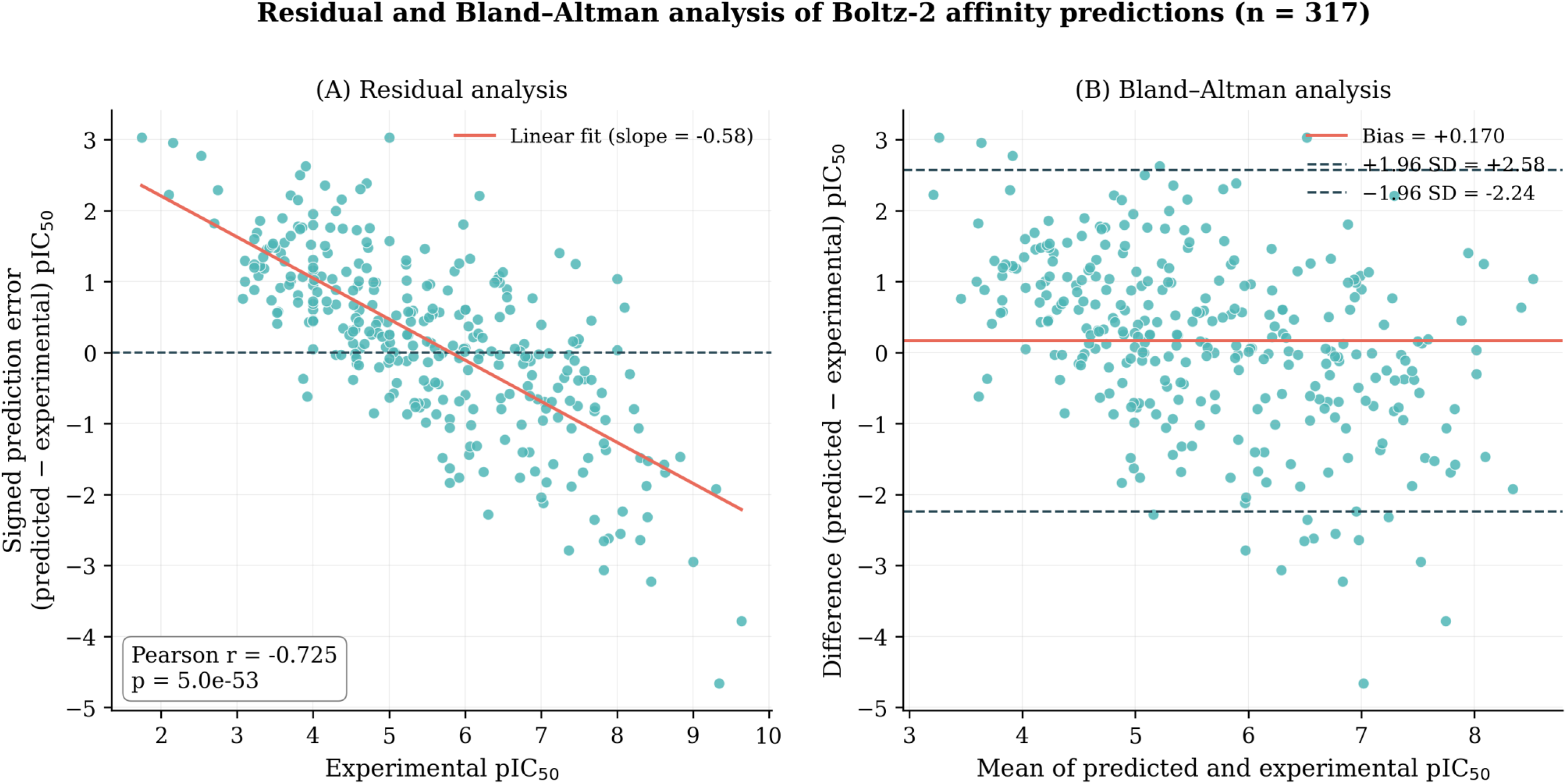
Residual and Bland-Altman analysis of affinity prediction error. (A) Signed prediction error (predicted minus experimental pIC50) plotted against experimental pIC50 for all 317 benchmark entries, illustrating the relationship between prediction bias and experimental potency. (B) Bland-Altman plot of the same data, showing the mean of predicted and experimental pIC50 (x-axis) against their signed difference (y-axis); the solid horizontal line indicates mean bias, and dashed lines indicate the 95% limits of agreement (bias ± 1.96 × SD of differences).

Bland-Altman analysis (Figure 7B) of the same 317 entries showed a mean bias of +0.170 pIC50 units (predicted minus experimental) with a standard deviation of differences of 1.227 units, yielding 95% limits of agreement of −2.24 to +2.58 pIC50 units.

### 5.6 Stratified Benchmark Performance

By protein target class. Using keyword-based classification of the associated protein name, the largest represented classes were kinases (n = 62), receptors (n = 36), phosphatases (n = 9), and proteases (n = 8), with the remaining 179 entries not matching a defined class (’Other’). Performance varied across these groups (Table 6): kinases, r = 0.54, MAE = 1.04; receptors, r = 0.78, MAE = 0.69; phosphatases, r = 0.64, MAE = 0.91; proteases, r = −0.41, MAE = 1.50 (n = 8); unclassified entries, r = 0.46, MAE = 1.01. The protease group’s correlation estimate is based on a small sample (n = 8) and is reported with that caveat.

By protein sequence length. Dividing the dataset into equal-sized tertiles by protein length, Pearson correlation was r = 0.54 (MAE = 1.00) for the shortest tertile (n = 106), r = 0.71 (MAE = 0.95) for the middle tertile (n = 105), and r = 0.52 (MAE = 0.95) for the longest tertile (n = 106).

By ligand size. Dividing the dataset into equal-sized tertiles by ligand heavy-atom count, Pearson correlation was r = 0.47 (MAE = 0.95) for the smallest tertile (n = 112), r = 0.65 (MAE = 0.93) for the middle tertile (n = 109), and r = 0.57 (MAE = 1.04) for the largest tertile (n = 96).

### 5.7 Representative Case Studies

Three entries illustrating the range of prediction agreement observed in the benchmark are presented in Figure 8.

**Figure 8.**
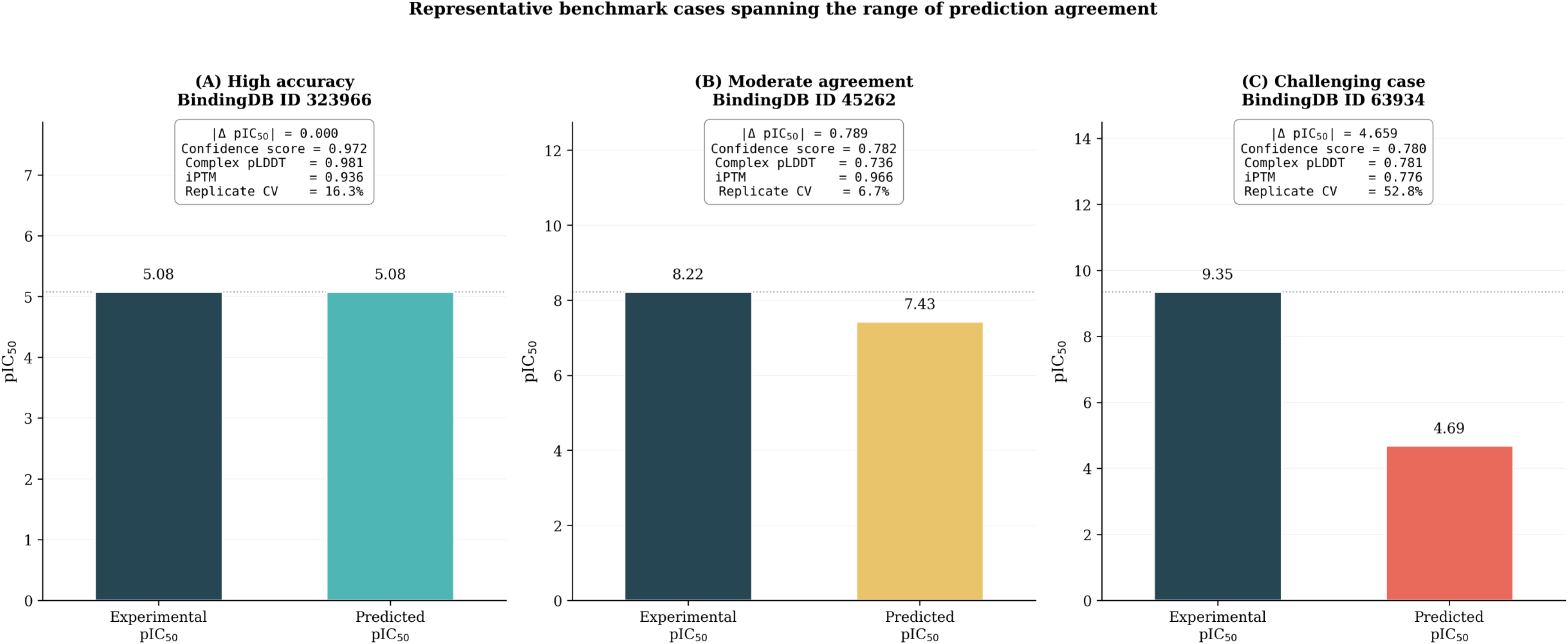
Representative case studies spanning the range of prediction agreement. Predicted structure, confidence metrics, and affinity values for three representative benchmark entries: (A) a high-accuracy prediction (BindingDB ID 323966; absolute error 0.0004 pIC50 units); (B) a moderate-agreement prediction (BindingDB ID 45262; absolute error 0.789 pIC50 units, approximating the dataset median); and (C) a challenging case with substantial underprediction (BindingDB ID 63934; absolute error 4.659 pIC50 units, the largest observed in the benchmark). Each panel reports experimental and predicted pIC50, confidence score, complex pLDDT, interface predicted TM-score (iPTM), and replicate coefficient of variation.

High accuracy. BindingDB entry 323966 (experimental pIC50 = 5.076) had a replicate-averaged predicted pIC50 of 5.076, an absolute error of 0.0004 pIC50 units - the smallest observed in the dataset. This entry had a confidence score of 0.972, complex pLDDT of 0.981, iPTM of 0.936, and a replicate coefficient of variation of 16.3%.

Moderate agreement. BindingDB entry 45262 (experimental pIC50 = 8.222) had a predicted pIC50 of 7.433, an absolute error of 0.789 pIC50 units, closely matching the dataset median absolute error (0.789 pIC50 units). This entry had a confidence score of 0.782, complex pLDDT of 0.736, iPTM of 0.966, and a replicate coefficient of variation of 6.7%.

Challenging case. BindingDB entry 63934 (experimental pIC50 = 9.347, among the highest-potency compounds in the dataset) had a predicted pIC50 of 4.688, an absolute error of 4.659 pIC50 units - the largest observed in the dataset. This entry had a confidence score of 0.780, complex pLDDT of 0.781, and iPTM of 0.776 - each within the range typical of the dataset as a whole (Section 5.4) - and a replicate coefficient of variation of 52.8%, higher than the dataset mean (31.4%) and median (25.7%).

## 6. Discussion

### 6.1 Principal Findings

This work makes two related but distinct contributions. First, we developed Boltz2-Notebook, a Colab-native interface that removes the local-GPU, CUDA, command-line, and manual-YAML requirements from Boltz-2 use, replacing them with interactive parameter construction, automated execution, and integrated confidence and affinity visualization within a single browser-based workflow, extended by a manifest-driven batch mode for multi-target screening. Second, we conducted an independent evaluation of Boltz-2’s affinity prediction performance on a newly curated, 317-pair BindingDB benchmark, generated and executed using the same input schema and execution logic the notebook automates, but run directly through the Boltz-2 command-line interface rather than through the notebook itself. The benchmark showed moderate agreement between predicted and experimental pIC50 (Pearson r = 0.609, R^2^ = 0.371, MAE = 0.968 pIC50 units), high triplicate reproducibility (pairwise replicate r = 0.97), a systematic compression of predicted values relative to the experimental dynamic range, and no measurable relationship between Boltz-2’s self-reported confidence metrics and the accuracy of its affinity predictions. Taken together, these findings characterize both the practical usability of the software described in this work and a specific, previously undocumented limitation of the affinity-prediction confidence outputs it exposes to users.

### 6.2 Comparison with Existing Platforms

Boltz2-Notebook’s relationship to the Boltz-2 command-line tool is analogous to the relationship ColabFold (Mirdita et al., 2022) established with AlphaFold2 and RoseTTAFold: neither modifies the underlying model, and both instead remove the local-installation and command-line barrier through Colab-hosted execution. This comparison is instructive with respect to scope as well as intent - ColabFold’s principal engineering contribution was accelerating multiple sequence alignment generation via MMseqs2 to make browser-based execution practical at scale, whereas Boltz-2’s affinity-and constraint-aware input schema is comparatively more complex than AlphaFold2’s sequence-only input, such that the corresponding accessibility problem for Boltz-2 is concentrated less in alignment generation and more in structured input specification, which is where Boltz2-Notebook’s parameter-generation layer is concentrated.

DeepMind’s AlphaFold Server offers a directly comparable accessibility model for AlphaFold3 (Abramson et al., 2024) - free, browser-based, no local installation - but differs from Boltz2-Notebook in scope: it restricts ligand input to a curated, non-custom library, does not report binding affinity, and its terms of service explicitly prohibit use ‘in connection with any automated system that predicts the binding or interaction of the protein with ligands or peptides’ (Google, 2024), precluding the batch, screening-style use supported by the batch notebook described in Section 3.6. Boltz2-Notebook’s feature set - custom SMILES ligands, affinity prediction, user-defined constraints, and batch execution - is accordingly better suited to small-molecule-focused, screening-oriented use cases than the AlphaFold Server, though this reflects a difference in the models each interface wraps and the terms under which each was released, rather than a general claim of superiority.

OpenFold (Ahdritz et al., 2024) serves a different purpose from either of the above and is a useful contrast case rather than a direct competitor: it is an open, retrained reimplementation of AlphaFold2 oriented toward reproducibility, trainability, and transparency of the modeling pipeline itself, not toward removing computational barriers for end users running inference. Boltz2-Notebook does not address model reproducibility or retraining in this sense; its contribution is confined to accessibility and workflow automation around a fixed, pretrained model.

The wider structural biology landscape also includes model architectures and complementary tools that are not accessibility solutions in the sense discussed above, but that clarify where Boltz-2 and Boltz2-Notebook sit methodologically. Baek et al.’s RoseTTAFold (Baek et al., 2021) and its all-atom extension developed by Krishna et al. (Krishna et al., 2024) established the underlying co-folding representations this generation of models builds on, but neither provides a Colab-native interface or affinity prediction. Diffusion-based docking methods such as DiffDock, introduced by Corso et al. (Corso et al., 2023), address a related but distinct problem: predicting a ligand pose against a fixed, typically experimentally determined receptor structure, rather than jointly co-folding the complex and estimating affinity as Boltz-2 does; DiffDock is accordingly a candidate baseline for future comparative benchmarking (Section 6.5) rather than a competing accessibility platform. Table 7 summarizes the comparison across Boltz-2, AlphaFold3, the AlphaFold Server, ColabFold, OpenFold, and Boltz2-Notebook.

Relative to the base Boltz-2 CLI, the specific engineering contributions of Boltz2-Notebook are the interactive YAML-schema compiler (Section 2.3), which constructs syntactically and structurally valid input from discrete, validated fields rather than requiring hand-authored configuration, and the manifest-driven batch execution engine (Section 2.6), which provides preflight validation, resumable queued processing, and ranked result aggregation not present in the base CLI. Automated confidence/affinity visualization and Drive-based output organization are useful but comparatively modest engineering conveniences rather than novel capabilities; the DNA/RNA, multi-entity, constraint, and affinity modeling capabilities themselves are inherited unmodified from Boltz-2 (Passaro et al., 2025) and are not claims made by this work.

### 6.3 Scientific Significance of the Benchmark

The correlation seen in this benchmark overall (r = 0.609, R2 = 0.371) suggests a moderate, but statistically robust, correlation between the predicted and experimentally measured affinities from Boltz-2 on a dataset curated independently. On an overall basis, this is in the same ballpark as the range of performance seen by Boltz-2’s original description (Passaro et al., 2025) for its own internal benchmarks, but the datasets, filtering criteria and evaluation conditions are sufficiently different that a direct numerical comparison is not appropriate. Here we show high triplicate reproducibility (pairwise r=0.97), implying that Boltz-2’s stochastic sampling procedure generates stable affinity estimates for a given input under repeated inference. This is a useful practical property that is independent of accuracy against experimental data, as a model could in principle be reproducible without being accurate, or accurate on average while being unstable for individual predictions.

We think the most important finding of this benchmark is the absence of a measurable correlation between the self-reported confidence measures of Boltz-2 and its real affinity prediction error. Structural confidence metrics, such as pLDDT and interface predicted TM-score, were developed and validated primarily as indicators of structural model quality, and their extension to affinity-prediction confidence has not, to our knowledge, been independently and quantitatively assessed prior to this work. This mismatch between reported confidence and empirical accuracy is in line with general findings in machine learning that predictive confidence does not automatically follow correctness without explicit calibration, as shown by Guo et al. for modern neural network classifiers in general (Guo et al., 2017). The result reported here is not to say that Boltz-2 affinity predictions are unreliable in an absolute sense – correlation with experimental data (Section 5.2) is a different matter from confidence calibration – but it does show that on this benchmark a user cannot use confidence score, iPTM or complex pLDDT to tell which specific predictions are more or less trustworthy. This is directly relevant to any workflow where predictions are triaged or prioritised by reported confidence, including the confidence dashboard enabled by Boltz2-Notebook (Section 3.4). We emphasise that this is a specific, actionable limitation, and not a general statement about model quality.

The systematic compression of the predicted affinity range relative to the experimental range, and the associated drop in stratum-specific correlation after the dataset was split by potency (Section 5.6), are consistent with regression-toward-the-mean behaviour that is frequently observed in machine-learning affinity predictors, and indicate that the pooled correlation coefficient is affected by the wide dynamic range of the benchmark and should not be extrapolated to suggest comparable discriminative power within a narrower potency range, such as typically encountered during lead optimisation.

### 6.4 Practical Applications

Within these performance characteristics, Boltz2-Notebook’s principal practical value is in lowering the technical barrier to using Boltz-2 for tasks including exploratory protein-ligand complex modeling, early-stage prioritization of candidate compounds via the batch screening workflow, structural hypothesis generation for protein engineering or variant design, and modeling of multi-chain or nucleic-acid-containing assemblies, for researchers who lack local GPU infrastructure or command-line proficiency. Its accessibility characteristics are also relevant to graduate and workshop-based teaching contexts, where hands-on exposure to modern structure-prediction methods is often constrained by institutional computing access rather than by conceptual difficulty. The confidence-calibration finding in Section 6.3 is directly relevant to how such applications should be conducted in practice: affinity outputs are more appropriately used for coarse ranking or hypothesis generation within a batch of candidates than as a confidence-weighted filter for individual predictions, given the absence of a demonstrated relationship between confidence and accuracy in this benchmark.

### 6.5 Limitations

Several limitations should be considered when interpreting this work. The benchmark comprises 317 protein–ligand pairs from a single source (BindingDB) restricted to single-chain, single-PDB targets with a strict IC50-only filter; this scope, while enabling a tractable and reproducible curation pipeline, excludes multimeric targets, EC50/Kd-reported assays, and larger protein or ligand systems, and the resulting performance estimates should not be assumed to generalize beyond this scope without further evaluation. No comparator method (e.g., a docking-based approach such as DiffDock (Corso et al., 2023) or AutoDock Vina (Trott & Olson, 2010), a physics-based approach such as the FEP+ protocol (Wang et al., 2015), alternative machine-learning affinity models, or Boltz-1) was evaluated on the same dataset, which limits the extent to which the reported correlation and error metrics can be judged as strong or weak in absolute, rather than relative-to-prior-literature, terms. We did not perform an explicit temporal or identity-based leakage assessment against Boltz-2’s training corpus; because large-scale affinity models of this kind are typically trained on corpora that may overlap with public repositories such as BindingDB, the possibility of some degree of train–test overlap cannot be excluded, and the reported performance should accordingly be interpreted as an upper-bound-compatible, rather than definitively leakage-free, estimate. The benchmark evaluates the underlying Boltz-2 model via direct command-line execution rather than the notebook software described in this manuscript, and while this was a deliberate design choice to allow scalable, unattended execution (Section 4.1), it means the benchmark does not itself constitute a validation of Boltz2-Notebook’s parameter-generation or execution logic; that validation is confined to functional demonstration rather than systematic testing (Supplementary Table S5). Boltz2-Notebook’s execution environment is additionally dependent on Google Colab’s free-tier resource allocation, which is subject to session time limits, variable hardware assignment, and memory constraints that may affect larger or more complex prediction jobs and are not fully characterized in this work. Finally, both the software and the benchmark are inherently dependent on the underlying Boltz-2 model; any limitation intrinsic to that model, including those identified in Section 6.3, is necessarily inherited by any workflow built on top of it.

### 6.6 Future Directions

Several extensions exist to overcome the above limitations. Further, additional independently sourced datasets, EC50/Kd measurements, multimeric targets, and at least one comparator method would improve the external validity of the reported performance estimates. A further formal leakage assessment on the training data of Boltz-2 and a stratified evaluation across a larger, more systematically defined set of target classes would help to clarify the conditions under which the observed correlation and calibration findings hold. Future software directions include automated correctness testing of the parameter-generation logic against the full Boltz-2 constraint schema, expanded runtime and throughput characterisation of the batch pipeline at larger scale, and continued alignment with future Boltz releases as the underlying model evolves. Like other community tools like ColabFold (Mirdita et al., 2022), the long-term utility of Boltz2-Notebook will depend partly on continued maintenance and partly on contributions and feedback from its user base as it gains adoption.

Taken together, this work provides an accessible, reproducible interface to Boltz-2 and an independent, appropriately scoped assessment of its affinity-prediction performance, including a concrete limitation - the lack of confidence-accuracy calibration - that has direct implications for how Boltz-2-based workflows, including but not limited to this one, should be used in practice. The contribution is intentionally modest in its claims: Boltz2-Notebook does not extend Boltz-2’s modeling capability, and the benchmark does not establish state-of-the-art affinity prediction performance; rather, together they lower a specific practical barrier to use and provide external, reproducible evidence relevant to interpreting Boltz-2’s affinity outputs responsibly.

## 7. Conclusion

Boltz-2 offers a flexible, full-featured model for joint structure and binding affinity prediction, but its practical application is limited by local GPU and CUDA dependencies, command-line operation, and a rigid, hand-authored YAML input schema -- obstacles that mostly burden researchers without dedicated computational infrastructure or advanced scripting expertise. Boltz2-Notebook was developed to fill this gap in accessibility. It does not change or increase Boltz-2’s modelling capability, but instead provides a Colab-native interface that automates environment setup, converts structured user input into valid Boltz-2 configurations, handles prediction execution, and formats confidence, affinity, and structural output into an interpretable, reproducible form, with a batch-processing mode that extends this workflow to multi-target screening. Rather than new predictive methodology, these contributions are best viewed as advances in usability and workflow-engineering in an accessibility pattern established for earlier-generation structure predictors such as ColabFold (Mirdita et al., 2022).

In parallel to this software, we have performed an independent assessment of the performance of Boltz-2 in affinity prediction on a newly curated benchmark from BindingDB. This assessment showed moderate, reproducible concordance of predicted and experimental binding affinities, and a specific, practically relevant limitation: In this benchmark, the self-reported confidence metrics of Boltz-2 failed to distinguish more accurate predictions from less accurate ones. This observation is offered as external evidence relevant to the responsible interpretation of Boltz-2 affinity outputs, irrespective of the software interface used to generate them.

The two elements – an easy-to-use platform for running the state-of-the-art open biomolecular model, and an independent performance assessment – are intended to be complementary, to reduce the technical barrier to using the model, and to give researchers a realistic, externally validated basis for interpreting its affinity predictions. As foundation models for predicting biological structure and interaction continue to make rapid progress, tools that facilitate the use of these models without specialised infrastructure, along with independent evaluation of their outputs, will continue to be necessary complements to methodological progress itself. We plan to keep Boltz2-Notebook as an open-source project, including future Boltz releases, more testing and contributions from the community as they become available.

## Data Availability

The curated BindingDB benchmark dataset (Supplementary Table S1), including all per-replicate and consensus predicted affinity values, confidence metrics, and associated experimental data, is available in the Supplementary Material accompanying this article and at 10.5281/zenodo.22830828. Raw BindingDB source data are publicly available from BindingDB (https://www.bindingdb.org) under BindingDB’s terms of use. Structural prediction outputs, confidence/affinity JSON files, and pLDDT/PAE arrays generated during the benchmark are available at 10.5281/zenodo.22830828. All data curation, YAML generation, and result-aggregation scripts required to reproduce the benchmark dataset from the original BindingDB release are provided as Supplementary Data and in the associated code repository (see Code Availability).

## Code Availability

Boltz2-Notebook is freely available as open-source software under the MIT license. The source code, including all four pipeline-stage scripts (environment setup, parameter generation, prediction execution, and analysis) and the batch-processing notebook, is hosted at the project’s GitHub repository: https://github.com/AtharvaTilewale/boltz2-notebook & https://github.com/cxbl-gbu/. A companion project website (https://atharvatilewale.github.io/boltz2-notebook/) provides direct links to launch notebooks in Google Colab without local installation. The specific software version used to generate the results reported in this manuscript is archived and assigned a permanent identifier at 10.5281/zenodo.22830828. The underlying Boltz-2 model and its native command-line interface are separately available at https://github.com/jwohlwend/boltz under their own respective license, and are unmodified by this work.

## Author Contributions

AT: Conceptualization; Methodology; Software; Validation; Formal analysis; Investigation; Data curation; Visualization; Writing - original draft; Writing - review & editing.

DP: Conceptualization; Data curation; Supervision; Project administration; Formal analysis; Investigation; Writing - original draft; Writing - review & editing.

## Declaration of Generative AI and AI-assisted technologies in the writing process

During the preparation of this work the authors have used ChatGPT (GPT-5; https://chatgpt.com/) and Claude Sonnet 5 (Anthropic, San Francisco, CA, USA) in order to improve the readability and language of the manuscript. After using this tool/service, authors have reviewed and edited the content as needed and take full responsibility for the content of the published article.

## Acknowledgements

The authors thank the developers of Boltz-1 and Boltz-2 for releasing their models as open-source software, without which this work would not have been possible. The authors acknowledge the University of Edinburgh’s Eddie high-performance computing cluster for computational resources used in generating the benchmark dataset, and Google Colaboratory for providing the free-tier GPU infrastructure underlying the Boltz2-Notebook platform. The authors thank BindingDB for maintaining and providing open access to the curated protein-ligand affinity data used in this benchmark. The authors acknowledge the Gujarat State Biotechnology Mission (GSBTM), Department of Science & Technology (DST), Gujarat, for resources in a project grant (GSBTM/ JD(R&D)/610/20-21/351 and GSBTM/JD(R&D)/626/22-23/00018348). The authors further acknowledge the Gujarat Biotechnology University, Department of Science & Technology (DST), Government of Gujarat) for providing the necessary infrastructure facilities.

## Table Captions

**Table 1. Feature comparison across Boltz2-Notebook notebook variants.** Summary of functional capabilities available in the stable single-job notebook (V1), the advanced single-job notebook (V2), and the batch notebook, including supported entity types, structural constraints, execution modes, and output handling.

**Table 2. Biomolecular entity types and structural constraints supported via the parameter-generation interface.** Listing of input entity types (protein, DNA, RNA, small-molecule ligand) and structural constraint mechanisms available through Boltz2-Notebook’s parameter-generation stage (param_gen.py), with the corresponding Boltz-2 YAML schema element each interface element constructs. All listed capabilities originate from Boltz-2’s native input schema; Boltz2-Notebook provides guided, validated construction of the corresponding configuration.

**Table 3. Configurable execution parameters and run-profile presets.** Summary of user-configurable prediction settings exposed by the execution stage (Boltz_Run.py), including recycling steps, diffusion sampling steps, number of diffusion samples, MSA pairing strategy and depth, and optional physics-based potentials refinement, together with the fast/balanced/high-quality preset profiles.

**Table 4. Summary statistics of the curated benchmark dataset.** Descriptive statistics for the final 317-entry BindingDB benchmark dataset, including number of unique proteins and ligands, protein sequence length distribution (minimum, maximum, mean, median), ligand heavy-atom count distribution, and experimental pIC50 range, mean, and median.

**Table 5. Overall affinity-prediction performance metrics.** Pearson correlation coefficient, Spearman rank correlation coefficient, coefficient of determination (R^2^), mean absolute error (MAE), and root-mean-squared error (RMSE) comparing replicate-averaged predicted pIC50 to experimental pIC50 across the 317-entry benchmark, with 95% confidence intervals derived from 5,000 bootstrap resamples of the dataset.

**Table 6. Stratified benchmark performance by protein target class, protein length, and ligand size.** Pearson correlation coefficient and mean absolute error (MAE) computed separately within subgroups defined by (i) keyword-based protein target classification (kinase, receptor, phosphatase, protease, unclassified), (ii) protein sequence length tertile, and (iii) ligand heavy-atom count tertile. Sample size (n) is reported for each subgroup.

**Table 7. Comparative positioning of Boltz2-Notebook relative to related structure-and affinity-prediction platforms.** Qualitative comparison of Boltz-2 (command-line), AlphaFold3, the AlphaFold Server, ColabFold, OpenFold, and Boltz2-Notebook across access model, local GPU requirement, input interface, support for custom small-molecule ligands, binding-affinity prediction, structural constraint support, and batch/high-throughput screening capability, based on publicly documented features and terms of use of each platform as of manuscript preparation.

## Supplementary Information

### Supplementary Figures

Figure S1. Screenshots of the interactive parameter-generation interface illustrating input widgets for protein, ligand, DNA/RNA, and constraint specification.

Figure S2. Example rendered dashboard output from the analysis stage, showing 3D structure visualization with pLDDT overlay, PAE matrix, and affinity/confidence summary panel for a representative job.

Figure S3. Full distribution histograms of Boltz-2 confidence score, interface predicted TM-score, and complex pLDDT across the benchmark dataset.

Figure S4. Extended residual diagnostics, including histogram of signed prediction error and quantile-quantile plot of residuals.

### Supplementary Tables

Table S1. Complete 317-row benchmark dataset (BindingDB identifier, protein sequence, ligand SMILES, associated PDB identifier, experimental IC50/pIC50, per-replicate and consensus predicted affinity values, and all extracted confidence metrics).

Table S2. Per-replicate (n = 951 jobs) confidence, affinity, and structural-quality output values prior to consensus averaging.

Table S3. Coefficient-of-variation summary statistics for replicate predictions, per target.

Table S4. Full software-metadata table (name, license, repository, language, dependencies, supported OS/environment, contact information).

### Additional Supplementary Material

Supplementary Data 1. BindingDB curation and cleaning scripts used to derive the benchmark dataset.

Supplementary Data 2. Benchmark YAML generation and result-aggregation scripts.

Supplementary Data 3. HPC job submission scripts and full dependency/environment specification used for benchmark execution.

Supplementary Notebook 1. Example completed Boltz2-Notebook (V2) session for a representative protein-ligand complex.

Supplementary Notebook 2. Example completed batch notebook session for a small multi-target set.

Supplementary Document 1. Example Boltz-2 YAML configuration files generated by param_gen.py for each supported feature.

Supplementary Document 2. User documentation, including installation-free quick-start instructions and a feature/parameter reference.

