## Supplemental Figures for "Boltz2-Notebook: An Interactive Google Colab Platform for Diffusion-Based Biomolecular Structure and Binding Affinity Prediction using the Boltz2 model"

### Slide 1
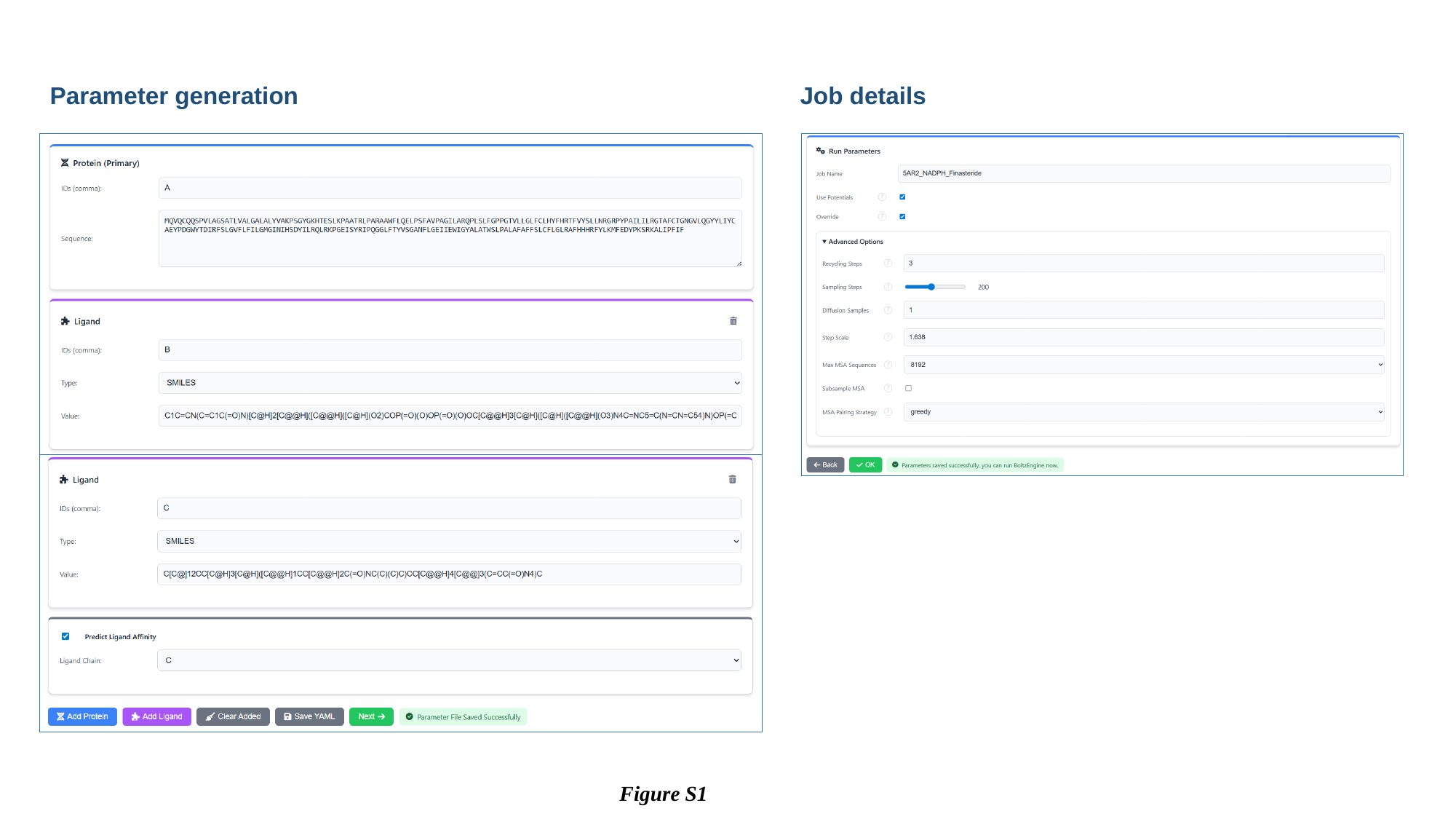

Parameter generation
Job details
Figure S1

### Slide 2
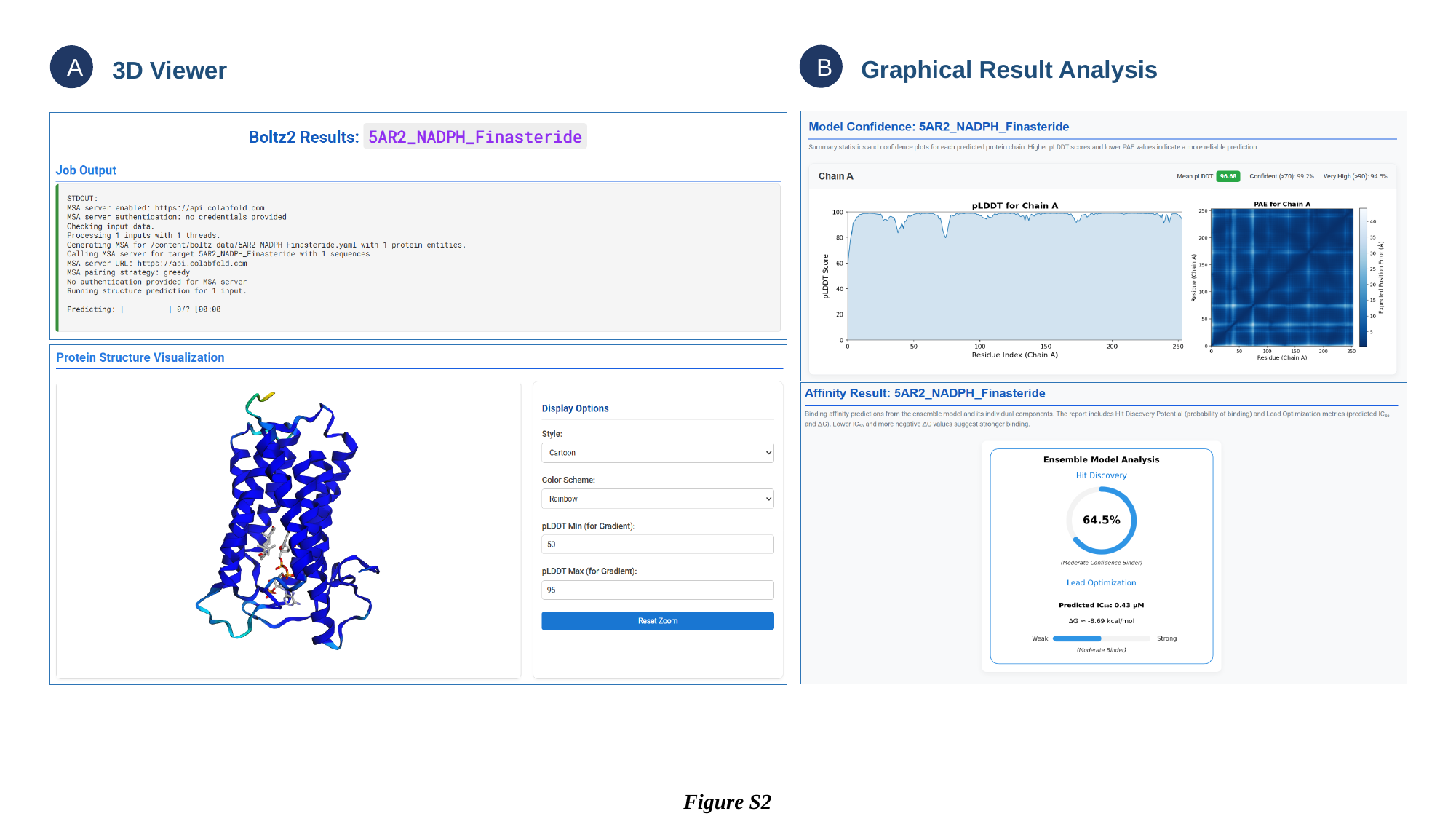

B
A
Graphical Result Analysis
3D Viewer
Figure S2

### Slide 3
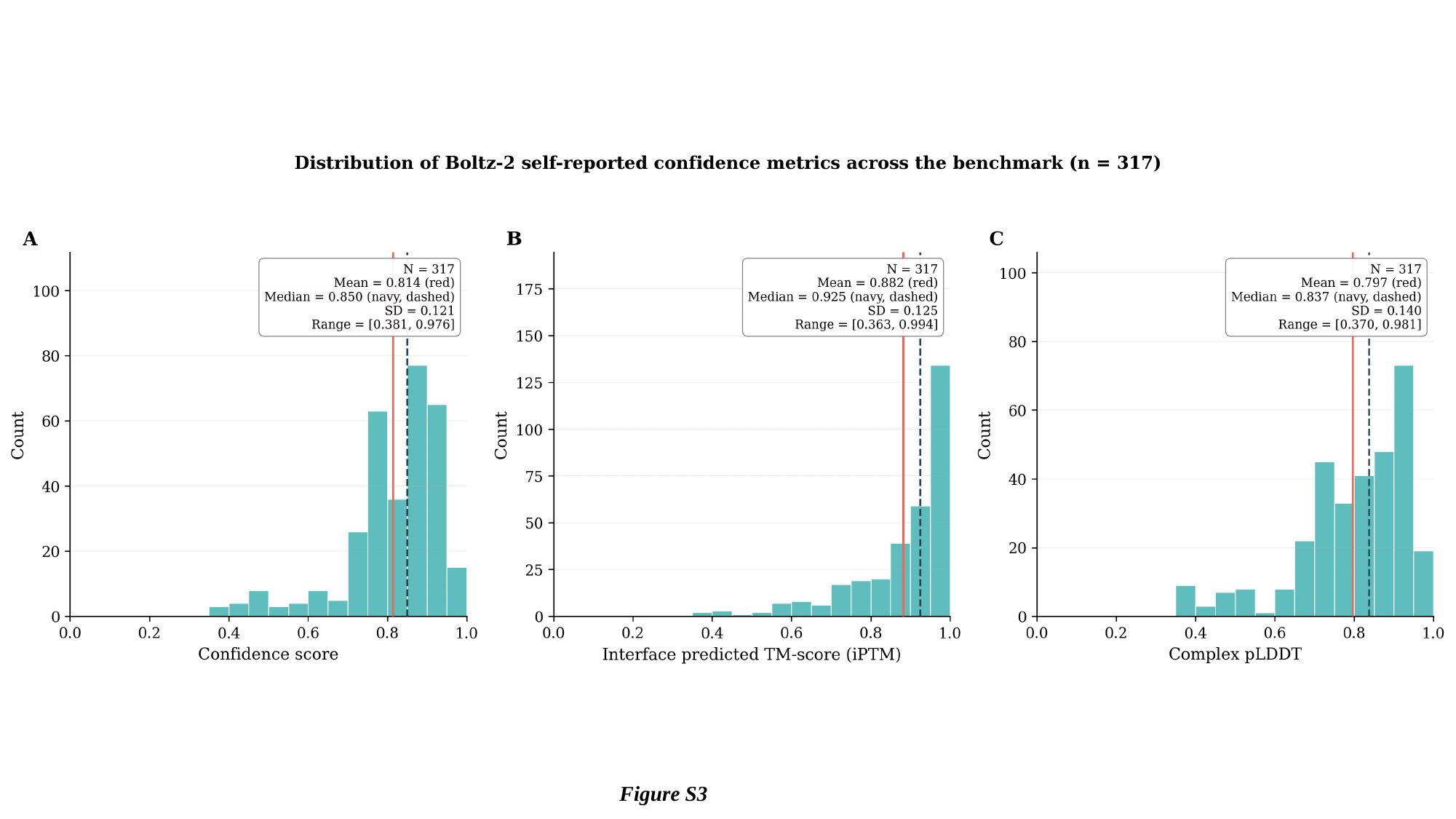

Figure S3

### Slide 4
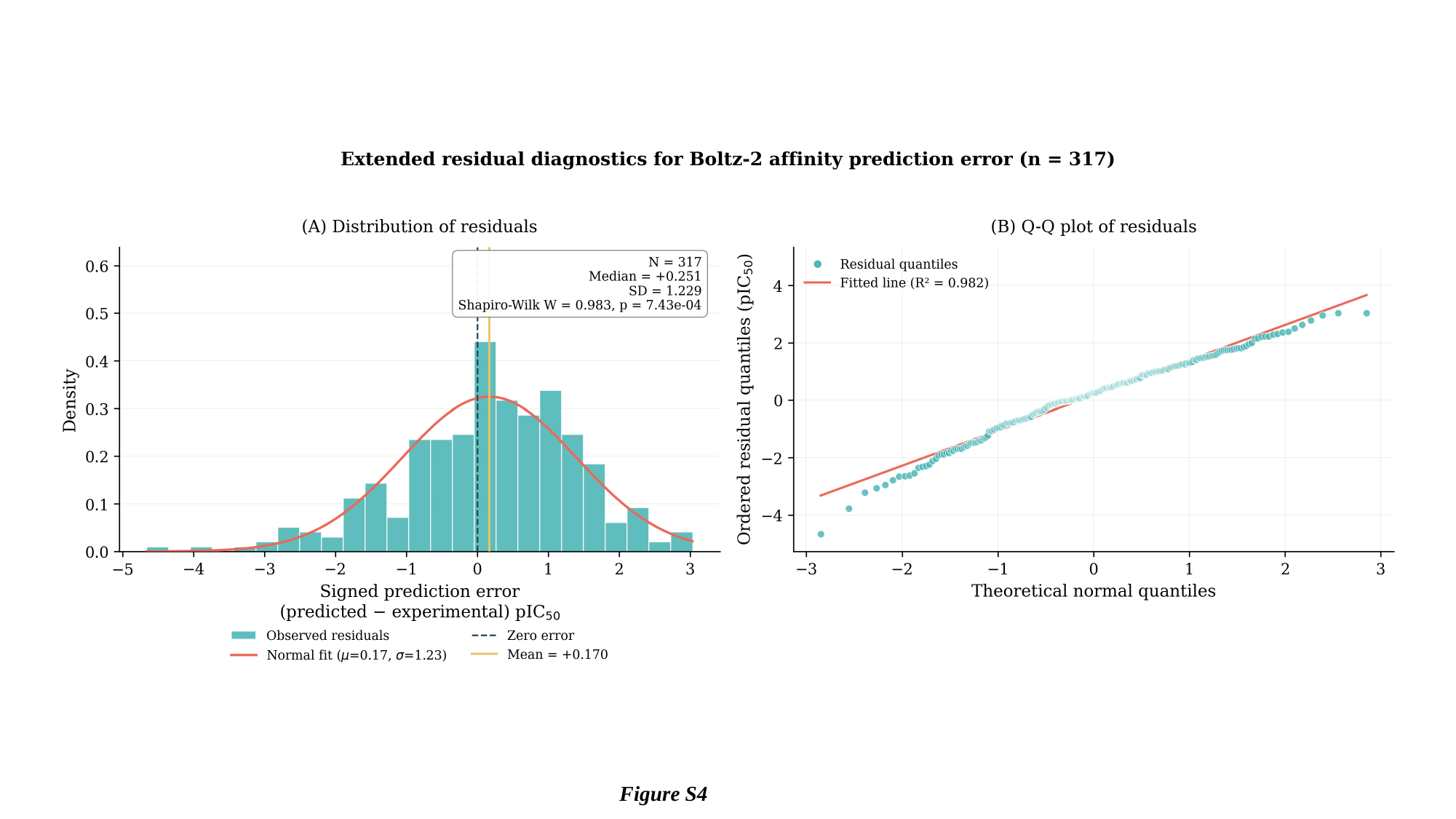

Figure S4
